# One-step generation of modified cattle and sheep from spermatid-like haploid stem cells

**DOI:** 10.1101/2024.09.10.612264

**Authors:** Lei Yang, Anqi Di, Lishuang Song, Xuefei Liu, Di Wu, Song Wang, Zhenting Hao, Lige Bu, Chunling Bai, Guanghua Su, Zhuying Wei, Li Zhang, Zhonghua Liu, Shaorong Gao, Guangpeng Li

## Abstract

Genome-modified ruminants, such as cattle and sheep, are important germplasm for agricultural applications. Haploid androgenetic stem cells (haSCs) can support the full-term development of embryos upon intracytoplasmic haSCs injection (iCHI) into intact oocytes. Thus, haSCs are invaluable resources for studying economic traits and significantly impact livestock breeding; however, ruminant haSCs have yet to be obtained. Here, we report the derivation of cattle and sheep haSCs using a recombinant FACE medium. These cells display characteristics of formative-state pluripotency and can differentiate into three germ layers, both *in vitro* and *in vivo*. Interestingly, the ectopic expression of protamine in haSCs converts their nuclei into a spermatid-like structure and synergistic enhancement of full-term development of iCHI embryos, named as Pro-iCHI approach. Furthermore, gene-modified cattle and sheep can be efficiently produced through the combinatorial use of the CRISPR and Pro-iCHI approach. Overall, Pro-iCHI provides an easy-to-manipulate agrobiotechnology tool for rapidly producing genetically modified livestock.

## Introduction

Cattle and sheep are important ruminants for the human food supply, including meat and milk, and are useful as experimental models for understanding developmental mechanisms and physiological functions. Efficient and precise genetic engineering in livestock holds great promise in agriculture breeding. Haploid androgenetic stem cells (haSCs) contain only one set of genomes inherited from the sperm and can support the production of viable transgenic rodents (mice and rats) through intracytoplasmic haSCs injection (iCHI) into oocytes ^1–3^. Electroporation-based or direct injection of the CRISPR/Cas system into zygotes have been established ^4, 5^, yet the emergence of haSCs not only offers a potential replacement for sperm but also presents an ideal strategy for loss-of-function screening assays ^6^. This is because a recessive mutation would yield a clear phenotype ^7^, as haSCs possess only one copy of each gene. Therefore, haSCs provide an avenue for genetic analysis from the cellular to the organismal level. However, livestock haSCs have not been achieved.

In this study, we applied the recombinant FACE medium for the derivation of bovine and ovine haSCs. These ruminant haSCs possess the ability to differentiate into various somatic cell types, as demonstrated through teratoma formation and chimeric assays. Furthermore, bovine haSCs can contribute to the germline of chimeric cattle and are competent for the *in vitro* induction of primordial germ cell-like cells (PGCLCs). Homogeneous gene editing in ruminant haSCs also can be efficiently achieved using the CRISPR-based prime editing system. Importantly, we found that the loss of asymmetrical histone modifications, specifically H3K4/K9/K27me3, in the pronuclei is a major barrier that impedes the development of iCHI embryos. This barrier can be overcome through the exogenous expression of protamine in ruminant haSCs, allowing both bovine and ovine iCHI embryos to develop to the blastocyst stage at a rate comparable to that of *in vitro* fertilization (IVF) and generating fertile cattle efficiently. Mechanistically, we showed that protamine alters the chromatin structure of ruminant haSCs, converting them into spermatid-like nuclei and removing the H3K4/K9/K27me3 modifications.

## Results

### Derivation of bovine and ovine haSCs using FACE medium

We first generated bovine haploid androgenetic embryos using two different methods: (i) injecting sperm into enucleated oocytes; (ii) removing the female pronucleus from zygotes (Fig. 1a, Extended Data Fig.1a). Expectedly, sperm actively underwent DNA demethylate in haploid embryos ^2^ (Extended Data Fig.1b). The blastocyst rate and quality of the injection and removal approaches were comparable (Fig. 1b,c, Extended Data Fig.1c,d, Supplementary Table 1), indicating that the sperm reprogramming by ooplasm was not affected by the manipulations. To generate bovine haSCs (b-haSCs), we adopted “naïve” or “primed” diploid SCs establishment mediums, which were originally developed for mice, humans, and cattle. The inner cell mass (ICM) was isolated from ha-blastocysts, plated on mitotically inactivated mouse embryonic fibroblast (MEF) feeder layers, and cultured in 9 different mediums (Extended Data Fig.2a). From 261 blastocysts, we only obtained outgrowths consisting of “master-regulators” OCT4^+^SOX2^+^ in LCDM, CTFR, and bEPSCM medium, but couldn’t establish stable cell lines (no more than eight passages; Extended Data Fig.2b-d).

**Fig. 1.**
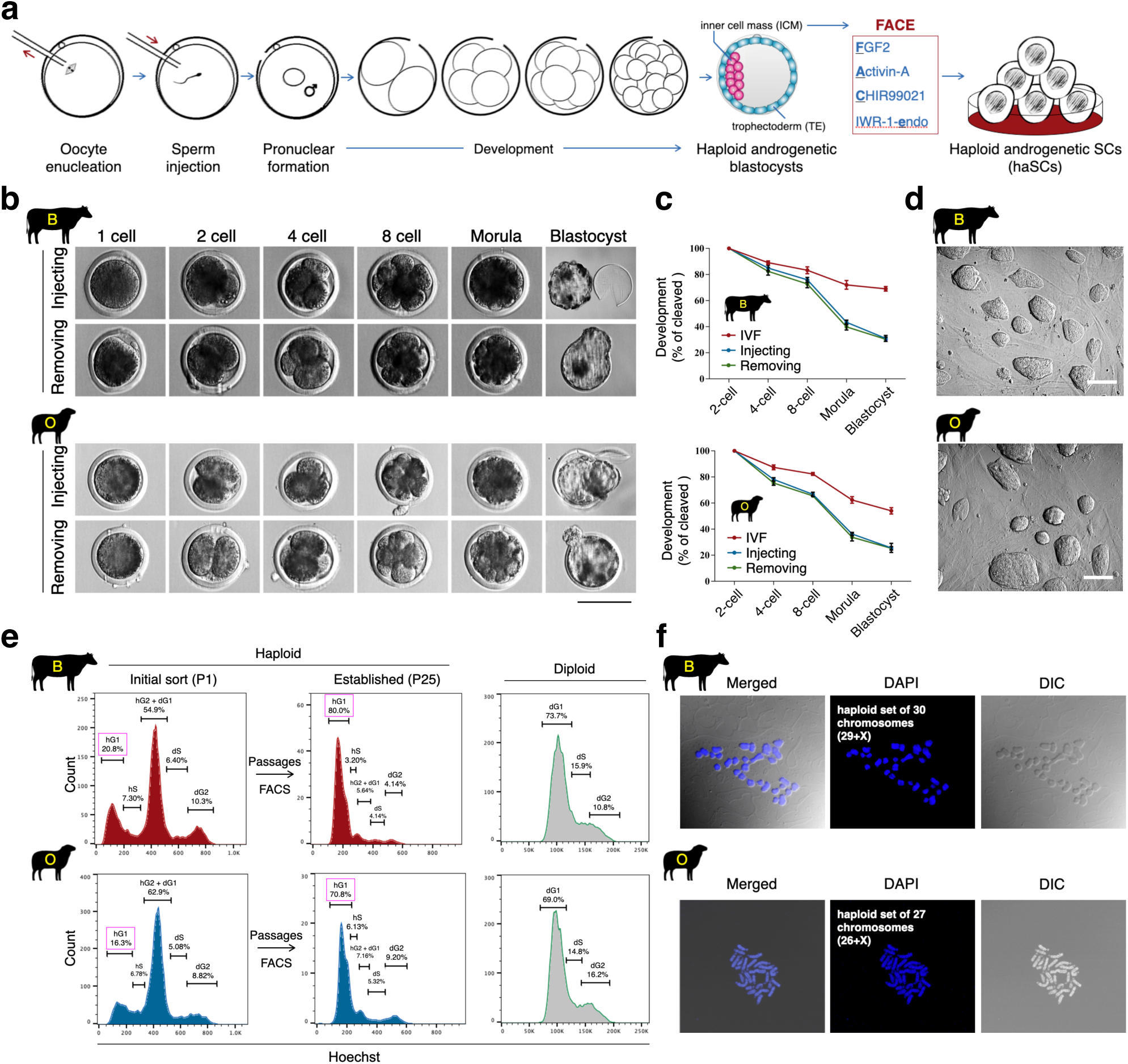
Efficient derivation of ruminant haSCs. a. Schematic overview of haSCs derivation. Haploid embryos were generated by injecting sperm into enucleated oocytes. ♂, male pronucleus. b. Representative images of ruminant androgenetic embryos. The haploid embryos are generated by injecting sperm into enucleated oocytes or removing the female pronucleus from normal diploid zygotes. Scale bar, 100 μm. c. Developmental efficiency of ruminant haploid and diploid embryos. Note that the haploid embryos generated through injecting sperm (31.16% in cattle, 25.68% in sheep) or by removing pronuclei (30.21% in cattle, 25.36% in sheep) showed no significant difference in the efficiency of blastocyst formation, while the blastocyst rate for haploid embryos was lower than that of diploid IVF embryos (68.84% in cattle, 53.71% in sheep). Shown is the percentage of embryos that reach the indicated stage. mean ± s.d., n > = 3 independent experiments. d. Morphology of ruminant haSCs derived from haploid ICMs. Scale bar, 100 μm. e. Effects of FACE medium on haploidy maintenance of ruminant haSCs with FACS enrichment. Diploid ruminant SCs were used as controls. f. Karyotype analysis of ruminant haSCs at passage 25. Bovine haploid chromosome set: 30 (29+X); Ovine haploid chromosome set: 27 (26+X).

To long-term maintenance of the b-haSCs, we screened different combinations of inhibitors and cytokines in the above successful culture medium (11 hits that target ten signal pathways; Extended Data Fig.3a). Cells cultured in FACE [FGF2::Activin-A::CHIR99021::IWR-1-endo] and FACX [FAC::XAV939] combinations formed colonies, while in others appeared differentiated or did not form colonies (Extended Data Fig.3b,c). Both IWR-1-endo and XAV939 act as Wnt/β-catenin inhibitors and can be used interchangeably ^8, 9^, thus [FACX] and [FACE] are essentially the same medium. We further titrated Activin-A and obtained cell lines that could be propagated for more than 80 passages in 10 ng/mL Activin-A, and haploid cells could be enriched by fluorescence-activated cell sorting (FACS; Fig. 1d-f, Extended Data Fig.2e, 3d, 3e, 4a-c).

We also succeeded in establishing ovine haSCs (o-haSCs) using FACE medium, which expressed pluripotency marks similar to that of b-haSCs (Fig. 1a-f, 2a, Extended Data Fig.1a-d, 2e, 4a-c, Supplementary Table 1). Consistent with rodent haSCs ^1–3^, no ruminant haSCs with the Y chromosome (Extended Data Fig.4a, c). Originally obtained b- and o-haSCs contain nearly 20% haploid cells, which was higher than that of murine haSCs ^1–3^ (< 5%; Fig. 1e). During gastrulation, the inner cell mass (ICM) of the blastocyst passes through three pluripotency states: naïve, formative, and primed ^10^ (Extended Data Fig.6a). Under the naïve state, mouse haSCs (m-haSCs) tend to rapidly diploidization ^2^, whereas the ruminant haSCs do not. These results prompted us to determine the pluripotency state of ruminant haSCs.

**Fig. 2.**
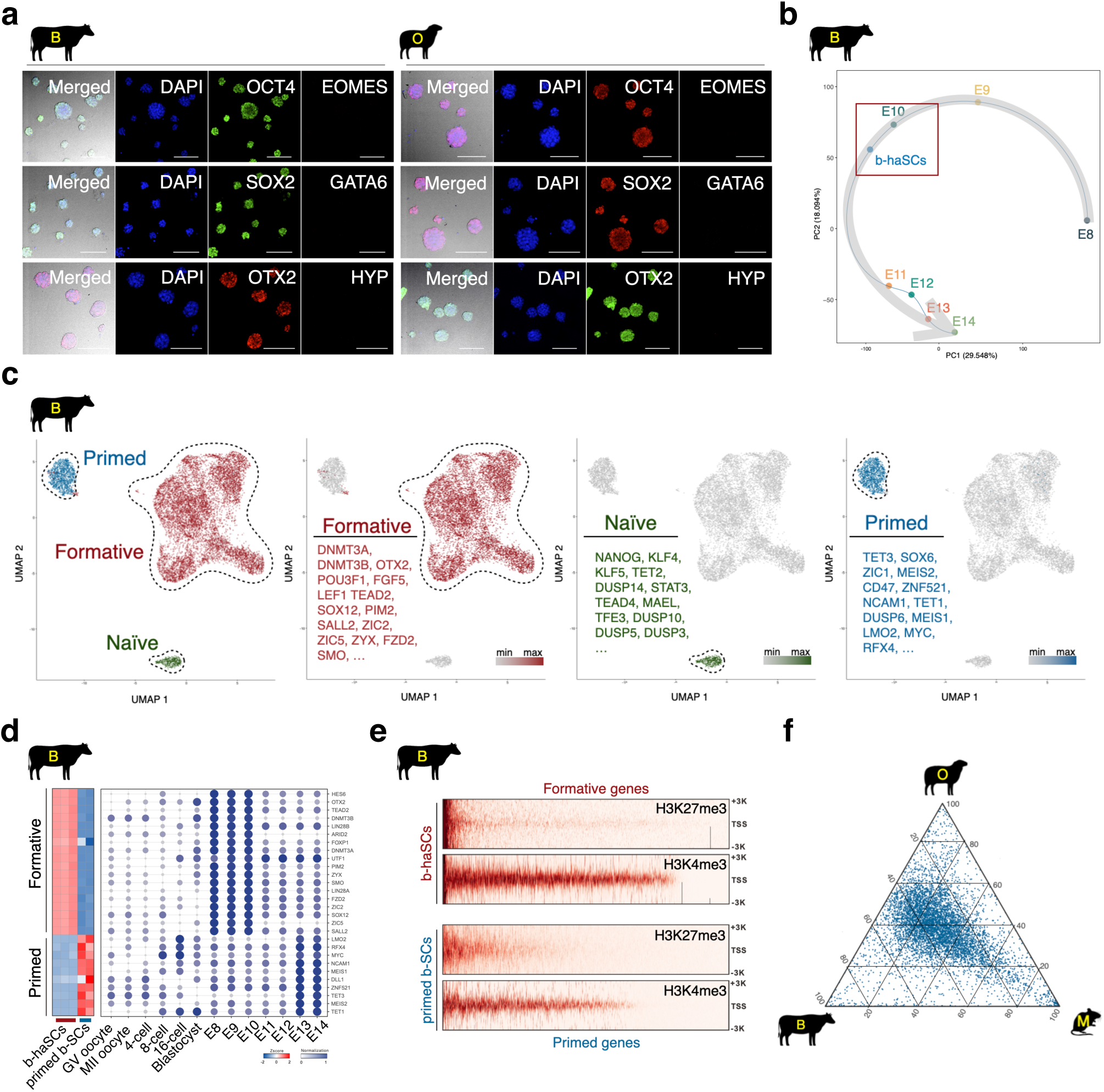
Molecular characterization of ruminant haSCs. a. Immunofluorescence images showing that ruminant haSCs expressed core pluripotency markers (SOX2 and OCT4) and formative pluripotency marker (OTX2). Scale bars, 25 μm. b. Scatter plot based on PCA of b-haSCs and bovine peri-implantation embryos (E8∼E14), showing the position of b-haSCs is closest to E10. c. UMAP projection of b-haSCs, coloured by pluripotency-associated marker genes. d. Heatmap and bubble displaying the expression of formative- and primed-pluripotency markers in different samples. e. Heatmap of H3K4me3 and H3K27me3 signals in b-haSCs and primed-b-SCs. f. Ternary plot showing cross-species comparison of orthologous transcription factors, formative pluripotency related genes.

### Pluripotency of bovine and ovine haSCs

We next analysed the transcriptome by bulk RNA sequencing (buRNA-seq) and found that the b-haSCs couldn’t belong to any stage of pre-implantation embryos ^11, 12^ (Extended Data Fig.5a,b). Meanwhile, b-haSCs also segregated from the recently reported diploid primed and expanded SCs ^13, 14^ (Extended Data Fig.5c,d), implying that b-haSCs are enriched in the features of formative pluripotency (Extended Data Fig.5e). To test this hypothesis, we performed single-cell (sc) RNA-seq during the gastrulation period of bovine epiblast from *in vivo* diploid embryos (E8∼14; Extended Data Fig.6a,b). Interestingly, b-haSCs were clustered with the embryos at E10 (Fig. 2b, Extended Data Fig.6b). We also identified three types of cells in b-haSCs: 83.8% formative, 12.1% primed, and a few 4.1% naïve (Fig. 2c,d, Extended Data Fig.7a,b).

To dissect the epigenetic features, we performed CUT&Tag assay (Extended Data Fig.8a). The H3K4me3 and H3K27me3 at the promoter of formative genes showed higher and lower in b-haSCs than primed b-SCs, respectively. Also, these modifications of primed genes exhibited the opposite trend (Fig. 2e, Extended Data Fig.8b). Consistently, the H3K4me3 and H3K27me3 cluster profiles segregated from primed b-SCs (Extended Data Fig.8c). The presence of H3K4me3 but not H3K27me3 in the promoter of basic-pluripotency genes, including OCT4, SOX2, and SALL4, is common to both b-haSCs and primed diploid b-SCs (Extended Data Fig.8d).

We also undertook a cross-species comparison, and found that o-haSCs clustered closer with b-haSCs than to primed o-SCs (Extended Data Fig.7c). Consistently, formative-related regulators were both present in bovine, ovine, and murine formative cells (Fig. 2f). The mRNA and protein of the formative markers, including DNMT3A, DNMT3B, and OTX2, were comparable among each species (Extended Data Fig.7d,e). We thus conclude that b- and o-haSCs harbour molecular features of formative pluripotency.

### Differentiation of bovine and ovine haSCs

We further tested the differentiation capacity of the ruminant haSCs through *in vitro* and *in vivo* assays. Expectedly, b- and o-haSCs could differentiate into three germ-line with around 16% haploid cells in embryoid bodies and teratomas, which is much higher than that of m-haSCs ^1^ (Fig. 3a,b, Extended Data Fig.10a-d, 11a-d). Both b- and o- haSCs could localize to the ICM positions of the intra-/inter-species chimeric blastocysts (Extended Data Fig.12a-d). Similar to teratoma formation, b- and o-haSCs possess the ability to differentiate into three germ layers in the intra-species chimaeras at E40 and E30, respectively, but no haploid cells were detected (Fig. 3c,d, Extended Data Fig.13a,b). Unlike o-haSCs, b-haSCs can contribute to the gonad (Extended Data Fig.13c,d). Consistently, b-haSCs are amenable to primordial germ cell-like cells (PGCLCs) induction *in vitro* (Fig. 3e,f; see Supplementary Note for more details). We also transferred inter-species chimeric blastocysts into foster mice, and detected b- haSCs giving rise to live inter-species chimaeras at E18.5 (Fig. 3g, Extended Data Fig.14a). However, the cattle-mouse chimaeras only reached 2-day-old (Extended Data Fig.14b-d).

**Fig. 3.**
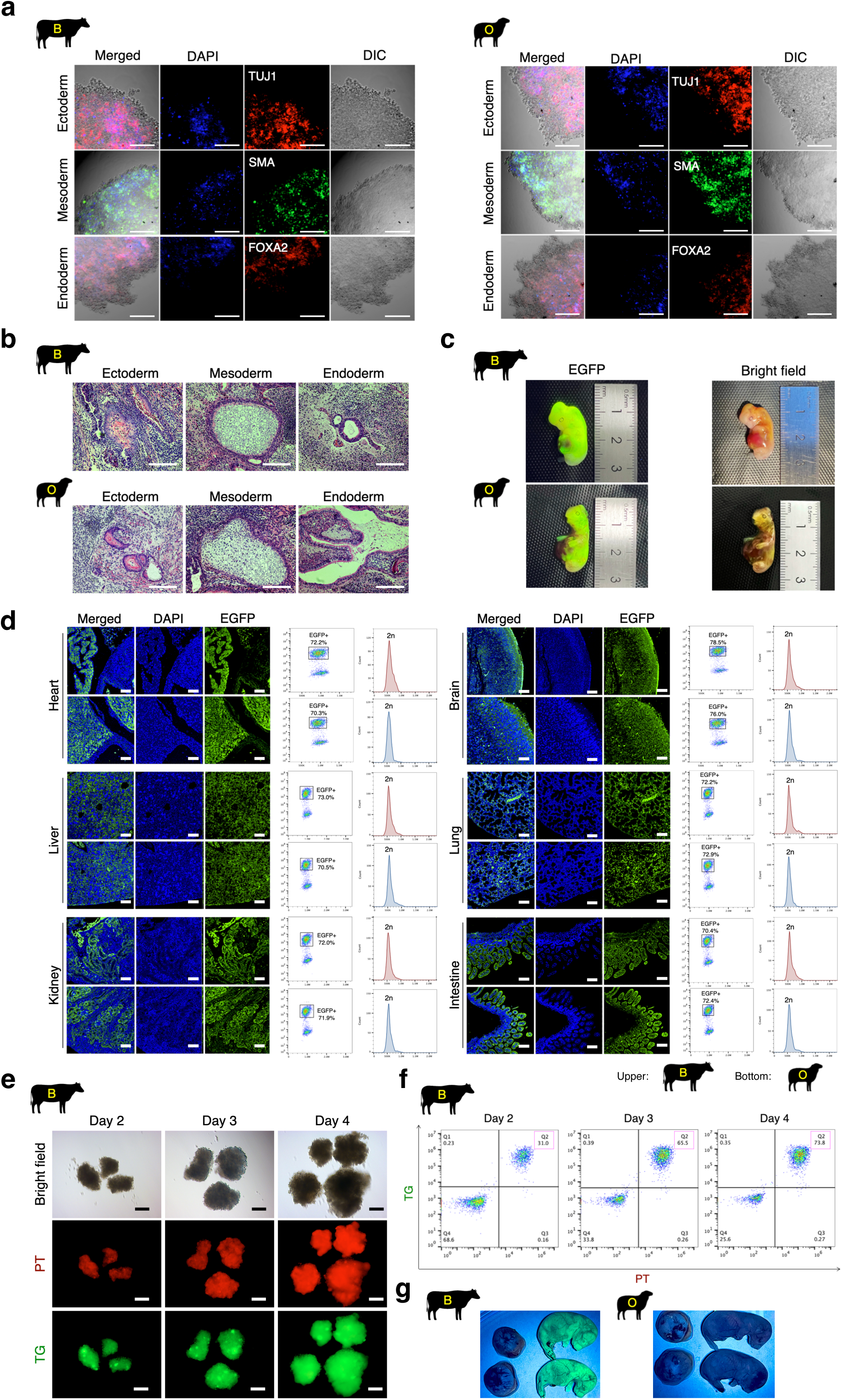
Functional differentiation analysis of ruminant haSCs. a. Immunostaining for three germ-line markers in embryoid bodies derived from ruminant haSCs, including ectoderm (TUJ1), mesoderm (SMA), and endoderm (FOXA2). Scale bar, 100 μm. b. Representative histochemical images of teratoma produced by ruminant haSCs. The dissection slices, including three germ layers, are stained with hematoxylin and eosin (H&E). Scale bar, 200 μm. c. Representative phase contrast and fluorescence images showing the chimeric contribution of EGFP-labeled b-haSCs or o-haSCs in the E40 cattle-cattle and E30 sheep-sheep chimaeras, respectively. d. Left panel: immunohistochemistry images showing EGFP-labeled b-haSCs or o-haSCs contributed to different tissues in the E40 cattle-cattle and E30 sheep-sheep chimaeras, respectively. Organ sections were stained with anti-GFP antibody. Right panel: FACS analysis of DNA content of the EGFP-positive cells within the different tissues. Scale bar, 200 μm. e. Representative phase contrast and fluorescence images showing PT::TG reporter signals during induction of PGCLCs from b-haSCs. PT, PRDM1/BLIMP1-tdTomato; TG, TFAP2C-mNeonGreen; Scale bar, 200 μm. f. FACS analysis of PT::TG reporter expression during induction of PGCLCs from b-haSCs. PT, PRDM1/BLIMP1-tdTomato; TG, TFAP2C-mNeonGreen. g. Inter-species chimeric cattle-mouse and sheep-mouse generated by EGFP-labeled ruminant haSCs at E18.5.

### Development of embryos reconstructed by bovine and ovine haSCs

We performed the iCHI experiment to replace sperm, by injecting the b- and o-haSCs into bovine and ovine oocytes, respectively ^2^ (Fig. 4a). The injected haSCs formed a pseudo-pronucleus and underwent DNA demethylation, which is consistent with *in vitro* fertilization (IVF) embryos (Fig. 4b, Extended Data Fig.15a,b). However, most b- and o-iCHI embryos could not develop to the blastocyst stage (Fig. 4c,d, Supplementary Table 2). We and other previous studies revealed that the aberrant methylation of DNA and histone is a barrier to the development of rodent iCHI and somatic cell nuclear transfer (SCNT) embryos ^15, 16^. We thus performed pyrosequencing and whole genome bisulfite sequencing (WGBS) on b- and o-haSCs, and detected that the differentially methylated regions (DMRs) of parental imprints maintained proper status as in sperm, including H19, IGF2, and DIO3 (Extended Data Fig.9a-c). These results suggest that DNA methylation does not cause the developmental failure of ruminant iCHI embryos.

**Fig. 4.**
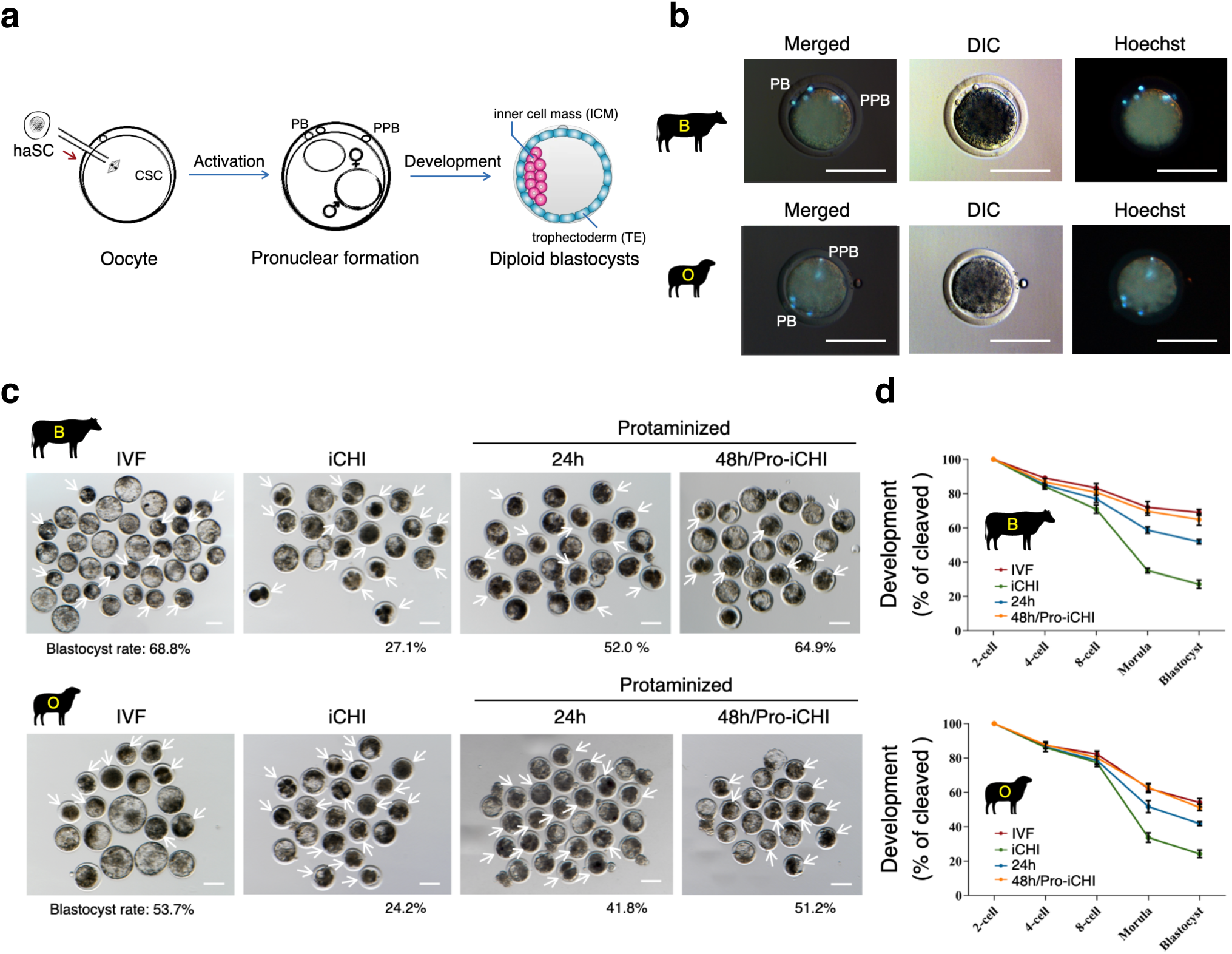
Evaluation of reconstructed embryos with ruminant haSCs. a. Schematic illustration for producing iCHI embryos. The metaphase ruminant haSCs were microinjected into matured MII oocytes. CSC, chromosome spindle complex; PB, polar body; PPB, pseudo-polar body; ♀, female pronucleus; ♂, male pseudo-pronucleus. b. Morphology of iCHI embryos at one-cell stage produced by ruminant haSCs. PB, polar body; PPB, pseudo-polar body; Scale bar, 100 μm. c. Morphology of preimplantation development of ruminant embryos at 7.5 days of *in vitro* culture. These embryos were derived from oocytes injected with wild-type or protaminized haSCs (24 and 48 hr post-protamine-induction). The IVF embryos were used as normal diploid control. Arrows indicate the degenerated embryos. Scale bar, 100 μm. d. Developmental efficiency of ruminant embryos. These embryos were derived from oocytes injected with wild-type or protaminized haSCs (24 and 48 hr post-protamine-induction). The IVF embryos were used as normal diploid control. mean ± s.d.; n > = 3 independent experiments.

### Generation of livestock by protaminized haSCs injection

Next, we analyzed the histone methylation. Unlike the asymmetric histone modification in IVF zygotes, H3K4/K9/K27me3 are present in all pronucleus of iCHI embryos at the one-cell stage (Fig. 5a, Extended Data Fig.16). However, injection with related demethylases KDM5B/4D/6A mRNA into iCHI embryos did not correct these errors (Extended Data Fig.17a-c, Supplementary Table 2). Moreover, KDM5B/4D/6A overexpression accelerated self-diploidization of ruminant haSCs (Extended Data Fig.17d). These results prompted us to find another method to rescue the development of iCHI embryos.

**Fig. 5.**
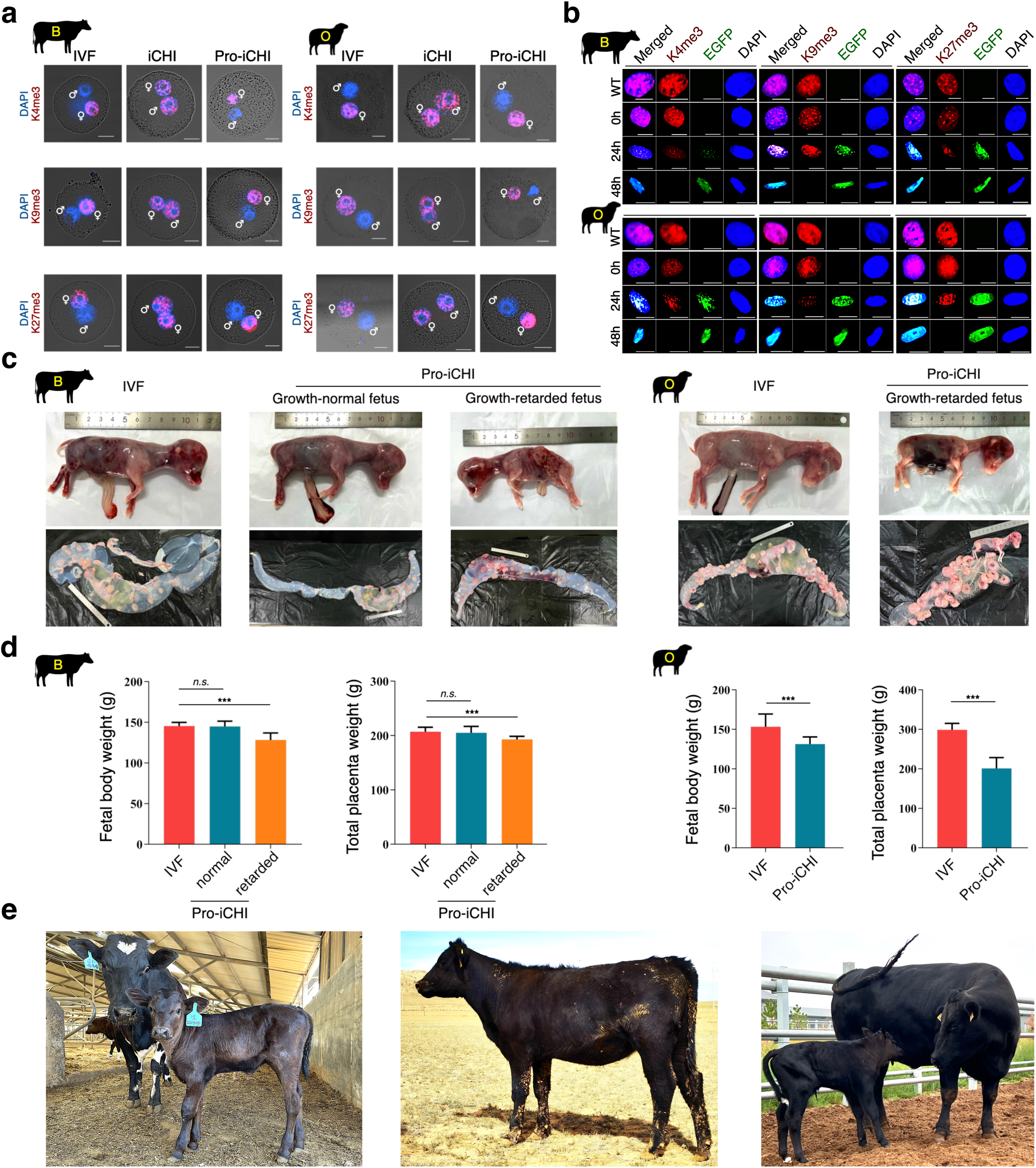
Protamine improves the development of reconstructed haSCs embryos. a. Immunostaining for H3K4me3, H3K9me3, and H3K27me3 in one-cell stage embryos derived by iCHI, Pro-iCHI, and IVF. ♀, female pronucleus; ♂, male pronucleus or pseudo-pronucleus; Scale bars, 25 μm. b. Immunostaining for H3K4me3, H3K9me3, and H3K27me3 in ruminant haSCs during protamine-induction (0, 24, and 48h post-protamine-induction; haSCs harbouring the dox-inducible PRM1-EGFP vector); WT, wild-type haSCs (untransfected with PRM1-EGFP); Scale bar, 20 μm. c. Representative images of ruminant conceptuses generated by Pro-iCHI or IVF, including the fetus proper (top panel), placenta, amniotic membrane, and allantoic membrane (bottom panel). The bovine and ovine conceptuses were collected at E75 and E65, respectively. The IVF conceptuses were used as normal control. d. Fetus and placenta weights of Pro-iCHI conceptuses, control groups were from IVF. The bovine and ovine conceptuses were collected at E75 and E65, respectively. ‘Normal’ Pro-iCHI fetus had a similar weight as the control IVF fetus, while ‘Retarded’ Pro-iCHI fetus had a smaller. mean ± s.d.; n = 3 independent experiments; n.s., not significant; \*\*\**P* < 0.001 by Student’s *t*-test. e. Representative images of Pro-iCHI cattle. Left, 1-month-old; Middle, 12-month-old; Right, 28-month-old and its offspring generated through natural mating with a wild-type male. Note that all the calves were naturally delivered by the surrogate cow.

As protamine (Pro) can exclude histone modifications during spermatogenesis ^17^, we used a non-integrated episomal vector to express the PRM1 gene transiently. During Pro-induction in b- and o-haSCs, we observed their nuclei progressively compacted into spermatid-like structures concomitant with H3K4/9/27me3 removal (Fig. 5b, Extended Data Fig.18a,b). The dynamics of “protaminization” were further confirmed by CUT&Tag assay in o-haSCs (Extended Data Fig.18c). Importantly, the symmetric errors observed in iCHI embryos can be amended by using 48-h protaminized haSCs as donors (Fig. 5a), which we refer to as Pro-iCHI approach. The blastocyst formation rate of embryos derived by Pro-iCHI was comparable to that of IVF-produced embryos (Fig. 4c,d, Extended Data Fig.16, Supplementary Table 2).

After embryo transfer, neither bovine (0/20) nor ovine (0/26) embryos produced by iCHI formed the conceptus (Extended Data Fig.19a, 20a; Supplementary Table 3). Contrarily, Pro-iCHI produced bovine and ovine embryos that could develop up to E75 (14/14) and E65 (17/20), respectively. Unlike SCNT animals with large offspring syndrome (LOS), bovine Pro-iCHI conceptus was either with normal IVF conceptus weights (2/14) or developmentally retarded (12/14; Fig. 5c,d, Extended Data Fig.19b, Supplementary Table 3), which is consistent with previous findings in rodents ^1–3^. Strikingly, we obtained one normal Pro-iCHI calf (14.3%, 1/7), and it developed into fertile adulthood (Fig. 5e, Supplementary Table 3). Additionally, the serum metabolomes of Pro-iCHI cattle are similar to that of IVF cattle (Extended Data Fig.21a-c). Unfortunately, all the ovine Pro-iCHI conceptus showed growth retardation and stillbirth (Fig. 5c,d, Extended Data Fig.20b,c, Supplementary Table 3).

Transcriptome analysis revealed that normal Pro-iCHI fetuses had similar expression patterns to IVF controls, whereas the expression of apoptosis and degeneration-related genes was upregulated in growth-retarded Pro-iCHI fetuses (Extended Data Fig.22a). We and others previously identified severe loss of H19 DMR in growth-retarded iCHI mice and rats ^3, 18^. Contrarily, we observed similar H19 methylation statuses between Pro-iCHI and IVF produced bovine and ovine fetuses, including normal and growth-retarded (Extended Data Fig.22b). Thus, further reduction of apoptosis and identification of other epigenetic barriers will provide clues for further improving the post-implantation development of ruminant Pro-iCHI embryos.

### Production of gene-edited animals from bovine and ovine haSCs

To evaluate the Pro-iCHI approach for producing gene-edited livestock, we selected the Myostatin (MSTN) gene to edit using CRISPR prime editing (PE) ^19^, as mutations in this gene lead to significantly increased muscle mass known as “double-muscling”^20^. Because the genomic integration of exogenous DNA increases biosafety risk, we developed an integration-free episomal-based PE (ePE) system. Indeed, ePE enables the production of nearly 100% short deletion (11 bp) and nucleotide substitution (G>A) in MSTN loci of the b- and o-haSCs, respectively (Fig. 6a,b, Extended Data Fig.23a-c). Finally, through the Pro-iCHI approach, we obtained two full-term MSTN-edited calves (13.3%, 2/15; survived to adulthood) and a much smaller MSTN-edited stillborn lamb (0.39%, 1/256; Fig. 6c,d, Extended Data Fig.24a-c, Supplementary Table 4). The imprinted region of H19 also maintained normal methylation status in these MSTN-edited lambs and calves (Extended Data Fig.22b).

**Fig. 6.**
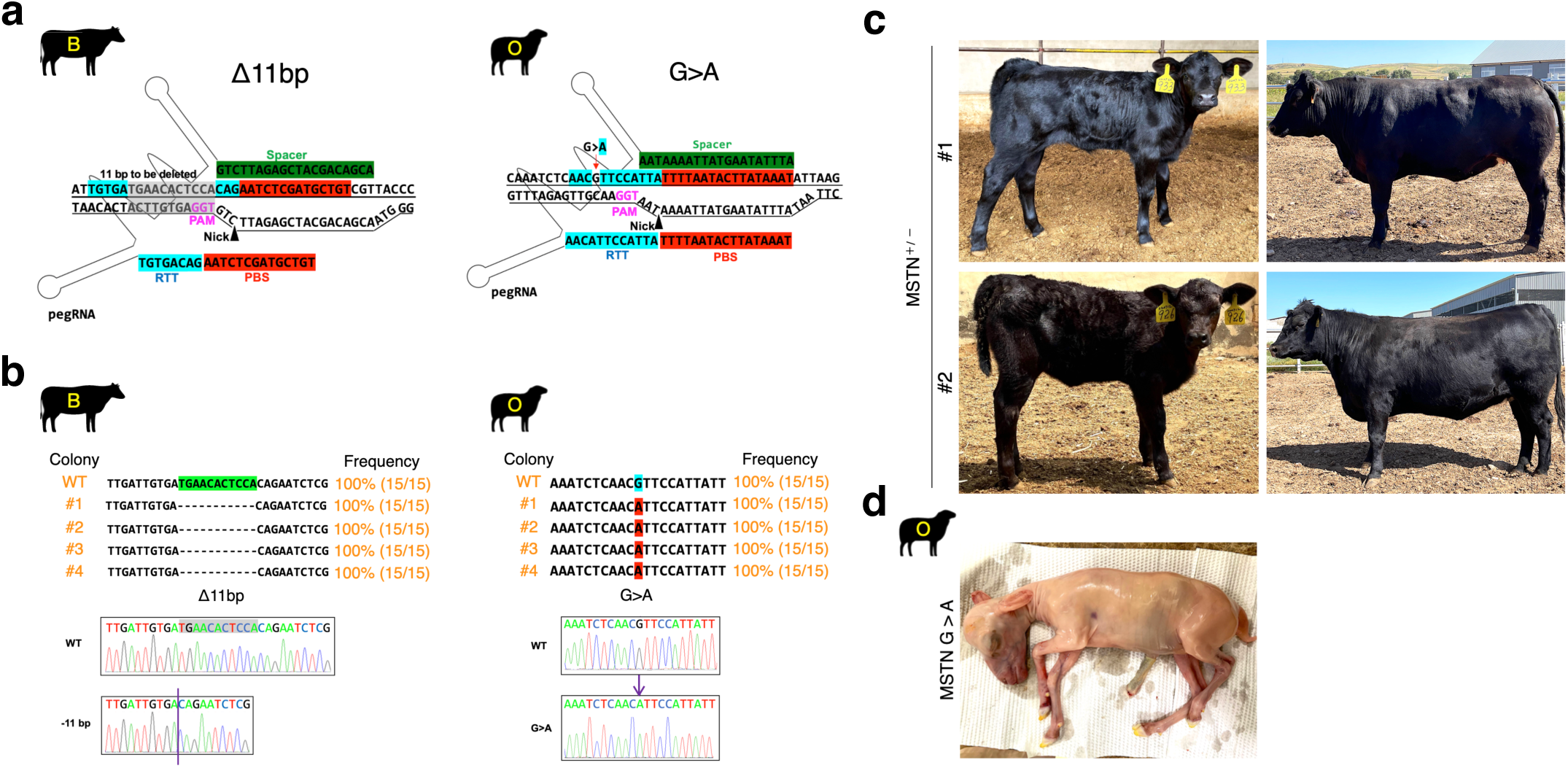
Application of ruminant haSCs in CRISPR studies. a. Schematic view of the MSTN-edit strategy via the CRISPR-ePE system. The pegRNA sequence and CRISPR proto-spacer adjacent motif (PAM) sequence are indicated. The MSTN loci was replaced with 11 bp deletion at exon 2 in b-haSCs and G>A substitution at 3ʹ-UTR in o-haSCs, respectively. b. Sanger sequencing results of the targeting site in ruminant haSCs. The efficiency of deletion (Δ) and point mutation (>) are indicated. PCR amplicons from the target regions in MSTN were analyzed using Sanger sequencing. Mutations were identified by aligning the sequenced amplicon to the wild-type (WT) MSTN sequence; Each line represents one individual haSCs sub-colony. Note that all the selected b- and o-haSCs sub-clones had an overt deletion or point mutation in the MSTN locus, with no WT sequence. c. Representative images of Pro-iCHI cattle derived from the MSTN-edited b-haSCs. Left, 1-month-old; Right: 12-month-old. Note that all the calves were naturally delivered by the surrogate cow. d. An image of Pro-iCHI sheep derived from MSTN-edited o-haSCs collected from the surrogate ewe after day 98 of embryo transfer.

## Discussion

In the past decade, we and others have successfully obtained haSCs from mice and rats using the 2i (PD0325901 and CHIR99021) medium ^1–3^, which help to maintain naïve-state pluripotency in rodents. However, ruminant haSCs cannot be established under the 2i conditions, consistent with earlier findings in diploid SCs, which showed that even low concentrations of the MEK1/2 inhibitor PD0325901 induced ruminant SCs death ^14^. Meanwhile, recently established the culture medium of primed or expanded pluripotency state for ruminant diploid SCs also cannot support the production of haSCs, including CTFR and bEPSCM medium ^13, 14^. Therefore, ruminant haSCs are more difficult to generate than their diploid counterparts, as well as rodent haSCs.

In this study, we performed an iterative small-molecule screening and identified a recombinant medium called FACE, which consists of FGF2, Activin-A, CHIR99021, and IWR-1-endo. This medium successfully supports the derivation of haSCs from both bovine and ovine haploid embryos. Unlike our previous observations of loss of paternal imprinting in rodent haSCs cultured in 2i medium ^1, 3, 18^, ruminant haSCs retain typical sperm-like imprinting under the FACE culture conditions.

Similar to the differentiation potential of diploid ruminant SCs and rodent haSCs ^1–3, 13, 14^, both bovine and ovine haSCs possess the ability to differentiate into various cell types of all three germ layers, as determined by the embryoid body formation *in vitro*, teratoma formation *in vivo*, as well as intra- and inter-species chimaera assays. While our results show that PGCLCs can be induced from bovine haSCs, further side-by-side functional comparisons between *in vitro* PGCLCs and *in vivo* PGCs still need to be performed. This not only provides an alternative source of gametes for livestock reproduction but also holds great promise for elucidating the basic mechanisms underlying human germ cell development. Because there are evident differences between the regulation of mouse and human in germline specification during their early post-implantation development ^21, 22, 23^.

Self-diploidization occurs rapidly during the differentiation of ruminant haSCs, which is consistent with our previous observation in mouse and rat haSCs ^1, 3^. Among the numerous studies to repress diploidization of mouse haSCs, the most simple method is directly adding the WEE1 inhibitor PD166285, CDK1 inhibitor RO3306, or ROCK inhibitor Y27632 to the rodent 2i medium ^24, 25^. It is possible that further optimization of the ruminant FACE medium by using the above small molecules may repress self-diploidization and improve the efficiency of haploid cell production.

With the combined advantages of haploidy and pluripotency in haSCs, precise editing in bovine and ovine genomes can be achieved through our developed integration-free ePE system. Despite the remarkable editing efficiency, iCHI embryos generated using neither bovine nor ovine haSCs failed to develop after implantation. We also uncovered that the loss of asymmetrical modifications of H3K4/K9/K27me3 at the one-cell stage is a major barrier to the development of ruminant iCHI embryos. Importantly, this developmental barrier can be overcome by simply using the protaminized haSCs as donor cells, enabling both bovine and ovine Pro-iCHI embryos to reach the blastocyst stage at an efficiency comparable to that of IVF. Consequently, the efficiency of the Pro-iCHI approach to produce both WT (1/7, 14.3%) and MSTN-edited cattle (13.3%, 2/15) was much higher, as compared to the conventional iCHI approach produced mice (29/660, 4.4%) and rat (6/1,056, 0.57%) ^1, 3^. During the revision process of this manuscript, two MSTN-edited cows produced by Pro-iCHI were pregnant for one month after mating with wild-type bulls.

Despite the significant improvement, more than half of the bovine and all the ovine embryos generated by the Pro-iCHI approach failed to develop after implantation. Nevertheless, the overall efficiency of live calf production is remarkable because it is higher than that of SCNT method performed by us and others ^26, 27^. To identify the other barriers for post-implantation development of Pro-iCHI embryos, we also isolated the IVF and Pro-iCHI produced bovine and ovine conceptus. Comparative morphological and transcriptomic analyses showed severe growth retardation and apoptosis in the abnormal Pro-iCHI embryos. Therefore, detailed analysis of the degeneration-related process will provide clues for further improving Pro-iCHI embryo development.

When we waited for the birth of Pro-iCHI cattle and their offspring (over 38 months), Smith and colleagues reported the isolation of diploid embryonic disc SCs (EDSCs) from ovine and bovine embryos using AFX medium ^28^. Our FACE medium contains the GSK-3 inhibitor CHIR99021, which enhances the self-renewal of mouse and human diploid SCs in combination with IWR-1-endo ^29^. Although XAV939 and IWR-1-endo are interchangeable inhibitors ^8, 9^, the concentrations of Activin-A (10 ng/mL in FACE vs. 20 ng/mL in AFX) and FGF2 (10 ng/mL in FACE vs. 12.5 ng/mL in AFX) in the above two medium also differ. Whether AFX medium can be used to generate haploid SCs with replaceable sperm competence warrants further investigation. Nonetheless, as a proof-of-concept, our study not only expands the usability of haSCs but also establishes protamine-assisted Pro-iCHI as an effective agrobiotechnology for producing trait-improved livestock.

## Methods

### Animals and chemicals

All animal procedures were performed under the ethical guidelines of the State Key Laboratory of Reproductive Regulation and Breeding of Grassland Livestock, College of Life Sciences, Inner Mongolia University (IMU). Specific pathogen-free-grade B6D2F1 (C57BL/6 × DBA/2), CD1, and Kun-Ming (KM) mice were purchased from Laboratory Animal Research Center (Inner Mongolia University) or Vital River Laboratories Co., Ltd. Livestock are raised at the experimental farm of IMU and processed in commercial slaughterhouses. Mice were housed in specific pathogen-free conditions with a 12-h dark/12-h light cycle. Unless otherwise indicated, all chemicals were purchased from Sigma-Aldrich Chemical Reagent Co., Ltd.

### Collection and *in vitro* maturation of oocytes

Collection and *in vitro* maturation (IVM) of ruminant oocytes followed the published protocols with slight modifications ^17, 30^. Briefly, bovine or ovine ovaries from the slaughterhouse were transported to the laboratory. Collect cumulus-oocyte complexes (COCs) and transfer them to the IVM medium for 24 h in 5% CO_2_ with humidified atmosphere at 38.5 ℃ [basic M199 medium supplemented with 2 mM glutamine, 300 μM, sodium pyruvate, 100 μM cysteamine, follicle-stimulating hormone (FSH; 50 ng/mL), luteinizing hormone (LH; 3 mg/mL), 1 μg/mL estradiol, and 0.04% gentamicin solution].

### Generation of haploid androgenetic embryos

Sperm of Angus cattle or Dorper sheep were prepared for manipulation according to the published protocols ^31–33^. The oocytes or zygotes were stained with 5 μg/mL Hoechst 33342 for 10 min to visualize spindle or pronuclei by a short 1-2 sec exposure to the ultraviolet (UV) light. Option A for injecting strategy, one-step SCNT was performed ^34, 35^, in which somatic cells were replaced by sperms. Briefly, the MII oocyte spindle was adjusted to 8 to 10 o’clock, and one donor sperm was injected into the nearby plasma. Immediately, the spindle was aspirated into the injection pipette and removed from the MII oocyte using the Piezo micromanipulator (Primetech) in the M2 medium containing 5 μg/mL cytochalasin B (CB) on a 37 °C heating stage of an inverted microscope (Nikon). After manipulation, the androgenetic embryos were activated in 5 μM ionomycin for 5 min and then incubated in 2 mM 6-dimethylaminopurine (6-DMAP) and 7.5 μg/mL CB in Ca^2+^-free KSOM-AA medium for 5 h. Option B for removing strategy, the zygotes were produced by *in vitro* fertilization (IVF) as previously described ^28^. Briefly, the zygotes were cultured in G1 medium for 7-8 h after IVF. Female pronuclei, which were near the second polar body, were removed from zygotes in the M2 medium containing 5 μg/mL CB using a blunt Piezo-driven pipette. Following activation or removing, the androgenetic embryos were cultured in G1 medium (Vitrolife) for 72 h, and then transferred into G2 medium (Vitrolife) for further culturing at 38.5 °C, humidity to saturation, 5% CO_2_, 5% O_2_, and 90% N_2_. The androgenetic embryos were allowed to develop into blastocysts that were used for haSCs derivation.

### Culture of ICMs using different SCs medium

The blastocysts were collected on day 8 post-activation. Only blastocysts scored as hatching (Stage 8) and excellence (Grade 1) with a well-developed inner cell mass (ICM), according to the Manual of the International Embryo Transfer Society (IETS), were selected for outgrowth culture. The SCs derivations were performed following the published protocols with slight modifications ^36^. In brief, blastocysts were treated with 1% actinase E (also known as pronase E) for 3-4 min under an inverted microscope with a warming stage [this treatment served to remove the zona pellucida (ZP) and loosen the cell junctions between ICM and trophectoderm (TE) cells]. Then, the blastocyst was held down using a hand-drawn glass Pasteur pipette with an internal diameter approximating the size of the ICM, and the ICM was aspirated into the pipette. The isolated ICMs were placed in individual wells of a 96-well dish seeded with a monolayer of feeder cells [the mitomycin C (MMC; 10 μg/mL for 2 h) treated mouse embryonic fibroblast (MEF) was used as the feeder, which was seed at a density of 20,000-25,000 cells/cm^2^ onto the pre-coated gelatin wells (0.1% gelatin for a minimum of 5 min at 37 °C)]. The ICMs were cultured with different mediums for SCs derivation, including 2i/L ^37^, 4i ^21^, t2iL+Gö ^38^, 5i/L/A ^39^, ABCL ^40^, LCDM ^41^, EPSC ^42, 43^, CTFR ^13, 44^, bEPSCM ^14^. After 10-14 days, following the mechanical transfer protocol ^45^, the outgrowths were cut into several clumps (150-200 cells) using the insulin needles (30-gauge, BD PrecisionGlide), and then replated in separate wells of a 12-well dish with feeder cells for further cultivation.

### Further culture of ICMs using different small-molecules

The ICMs and culture dishes were prepared using the same methods as described above. Individual ICMs were placed in separate wells of a 12-well dish seeded with a monolayer of MMC-treated MEFs overlaid with gelatin, and cultured in basal N2B27 ^46^ medium supplemented with 15% knockout serum replacement (KSR; Thermo), and different small-molecule compounds (Extended Data Fig.S3a). The small-molecules are used at the following concentrations: 3 μM CHIR99021 (C; Selleckchem), 2 μM Dimethindene maleate (D; Tocris), 2 μM Minocycline hydrochloride (M; Santacruz), 10 ng/mL LIF (L; Millipore), 2.5 μM IWR-1-endo (E; Selleckchem), 10 ng/mL FGF2 (F; Peprotech), 5 μM XAV939 (X; Selleckchem), 0.3 μM WH-4-023 (W; Selleckchem), 1 ng/mL TGFβ (T; Peprotech), 20 ng/mL Activin-A (A; Peprotech), 50 ng/mL BMP4 (B; R&D systems), 50 μg/mL Vitamin C (V; Selleckchem). All small-molecules are dissolved in dimethyl sulfoxide (DMSO). After 10-14 days, following the mechanical transfer protocol ^45^, the outgrowths were cut into several clumps (150-200 cells) using the insulin needles (30-gauge, BD PrecisionGlide), and then replated in separate wells of a 12-well dish with feeder cells for further cultivation.

### Derivation and routine culture of haSCs

The ICMs and culture dishes were prepared using the same methods as described above. Each ICM was plated into one well of a 96-well plate with the MMC-treated MEFs feeder in FACE medium at 37 °C, humidity to saturation, 5% CO_2_, 5% O_2_, and 90% N_2_. The medium was changed by half every 2 days. After 10-14 days, following the mechanical transfer protocol ^45^, the outgrowths were cut into several clumps (150-200 cells) using the insulin needles (30-gauge, BD PrecisionGlide), and then replated in separate wells of a 12-well dish with feeder cells. Cell expansion was carried out from 48-well to 6-well plates with feeders overlaid with gelatin and then to 6-well dishes for routine culture. Established haSCs were cultured on the MMC-treated MEFs feeders overlaid with gelatin in FACE medium, with daily media change. For passaging, the cells were disaggregated using TrypLE (Thermo) or Accutase (Stemcell) and passaged onto new feeder layers overlaid with gelatin at a split ratio of 1:10 every 5 days (TrpLE and Accutase have similar effects and can be used interchangeably). To enhance the survival of haSCs, 10 μM Y-27632 (ROCK inhibitor, Stemcell) was recommended to be added into the FACE medium for 24 h after the passage. The haploid genome integrity was maintained through FACS-enrichment of haploid cells every 15 passages.

The basal medium for FACE is mTeSR1 (Stemcell, based on Cat. No. 85850 medium without the addition of the 5X supplement medium) ^47^ or N2B27 ^46^ [total 500 mL was prepared as follows: 240 mL DMEM/F12 (Thermo), 240 mL neurobasal (Thermo), 2.5 mL N2 supplement (Thermo), 5 mL B27 supplement (Thermo), 1% GlutaMAX (Thermo), 1% nonessential amino acids (Thermo), 0.1 mM β-mercaptoethanol (Thermo), 1% penicillin-streptomycin (Thermo), and 15% KSR (Thermo)]. To prepare the FACE medium, small molecules and cytokines were added in the basal medium at the following final concentrations: 10 ng/mL FGF2 (F; Peprotech), 10 ng/mL Activin-A (A; Peprotech), 3 μM CHIR99021 (C; Selleckchem), and 2.5 μM IWR-1-endo (E; Selleckchem).

### FACS-enrichment of haploid cells

To sort haploid cells, we used fluorescence activating cell sorter (FACS) according to the published protocol with slight modifications ^48–50^. Briefly, haSCs were dissociated into single cells with TrypLE (Thermo) or Accutase (Stemcell) and then incubated with 15 μg/mL Hoechst 33342 for 15 min in a 37 °C water bath. Following centrifugation, cells were rinsed with Dulbecco’s phosphate-buffered saline (PBS) for removal of the Hoechst 33342, and resuspended in FACE medium with 10 μM Y-27632 (Stemcell). Subsequently, the stained cells were through 40 μm cell strainer and the haploid cells were sorted using FACSAria II (BD biosciences). After enrichment, the haploid cells were plated on fresh MEFs feeder in FACE medium containing 10 μM Y-27632. After 24 h, haploid cells were cultured in FACE medium without Y-27632. Usually, haSC clones were formed within 10 days. For cell analysis, after fixation in 70% ethanol, cells were digested by 20 mg/ml RNase A and stained with 50 mg/mL propidium iodide (PI). Then, the haploid cells were analyzed using CytoFLEX LX (Beckman). Analytic flow profiles were performed using FlowJo software (Ashland). FACS gating examples for haploid cells are shown in Supplementary Fig. 5, 6.

### PGC induction *in vitro*

Primordial germ cell (PGC) induction was performed as described previously ^51^. For the pre-culture stage, 3 × 10^5^ bES cells were placed in a 12-well dish coated with human plasma fibronectin (Millipore) and cultured for 24 h in NK5 medium: N2B27 medium containing 5% KSR (Thermo), supplemented with 100 ng/mL BMP4 (R&D systems), 6 μM CHIR99021 (Selleckchem), 2.5 μM IWR-1-endo (Selleckchem), and 10 μM Y-27632 (Stemcell) at 37.0 °C and 5% CO_2_. For the PGC induction stage, the above pre-induced cells were seeded into each well of low-binding U-bottom 96-well dish at a concentration of 6,000 cells/well, and cultured for 4 days in induction medium: GK15 medium containing GMEM medium (Thermo), 15% KSR, 0.1 mM nonessential amino acid (Thermo), 2 mM GlutaMAX (Thermo), 1 mM sodium pyruvate (Thermo), 0.1 mM β-mercaptoethanol (Thermo), 1% penicillin-streptomycin (Thermo), supplemented with 200 ng/mL BMP4, 100 ng/mL SCF (R&D systems), 50 ng/mL epidermal growth factor (EGF, Peprotech), 10 ng/mL LIF (Millipore), 6 μM CHIR99021, 2.5 μM IWR-1-endo, and 10 μM Y-27632 at 37.0 °C and 5% CO_2_.

### Immunofluorescence staining

Immunofluorescence (IF) was performed as described previously ^34^. Briefly, the cells were fixed with 4% paraformaldehyde (PFA) in PBS at RT for 1 h, and permeabilised in PBS supplemented with 0.2% Triton X-100 at RT for 45 min. The cells were incubated with primary antibodies in PBS that contained 3% blocker BSA (Thermo) at 4 °C overnight, then incubated with secondary antibodies for 1 h at RT in the dark. For 5mC and 5hmC staining, permeabilised embryos were incubated in 4 N HCl treatment for 10 min at RT, and then the oocytes were immediately transferred to Tris-HCl (pH 8.5) for 10 min to neutralise the residual HCl. Finally, the cells were mounted in the anti-fade solution with DAPI (Thermo) and compressed with a cover slip. All samples were observed by a laser scanning A1R microscope (Nikon). The images were analyzed using NIS-Elements C (Nikon). The antibody information is presented in Supplementary Table 5.

### Western blot

Western blot (WB) was performed as described previously ^16^. In brief, cells were lysed in RIPA lysis buffer (Beyotime) with protease and phosphatase inhibitor (Thermo). Whole-cell lysates were separated by sodium dodecyl sulfate-polyacrylamide gel electrophoresis (SDS-PAGE). Following electrophoresis (90 V for 30 min, 30-50 V overnight), the proteins were transferred from the gel to 0.2 mm PVDF membranes (Bio-sharp), and blocked for 2 h at RT in TBST buffer containing 5% BSA or 5% fat-free milk. Films were incubated with the primary antibodies in TBST buffer containing 3% BSA at 4 °C overnight. After washing three times in TBST, PVDF membranes were incubated with secondary antibodies for 1 h at RT. Then, bound antibodies were detected with the SuperSignal West Femto Substrate Trial Kit (Thermo). The antibody information is presented in Supplementary Table 5.

### Karyotype analysis

Before karyotype analysis, haSCs were incubated with 0.1 μg/mL KaryoMAX Colcemid (Thermo) in FACE medium for 3-4 h, and mechanically separated from the MEFs feeder using pipette tips. Then, haSCs were disaggregated into single cells by 0.05% trypsin-EDTA (Thermo) at 37 °C for 10 min. After centrifugation, the pellet was re-suspended in hypotonic solution (0.56% KCl) at RT for 10 min. Cells were then fixed with ice-cold fixative (3:1 methanol: acetic acid) at RT for 5 min three times. Finally, haSCs suspension was dropped onto an ice-cold fixative washed slide and stained with 4′,6-diamidino-2-phenylindole (DAPI; Thermo). For each cell line, more than 30 metaphases were examined.

### Chimeric assay

The chimeric assay was performed as previously described with slight modification ^34^. In brief, haSCs were used 1 day before passaging, which showed an optimal undifferentiated morphology. For interspecies chimaeras, fifteen haSCs labelled by EGFP fluorescence were microinjected into CD1 or KM mouse blastocysts using a blunt Piezo-driven pipette. Then, the reconstructed blastocysts were transplanted into the uterus of pseudo-pregnant mice at 3.5 days post-coitum (dpc). For intraspecies chimaeras, fifteen b- and o-haSCs labelled by EGFP fluorescence were microinjected into bovine and ovine blastocysts using a blunt Piezo-driven pipette, respectively. Then, the reconstructed embryos were transplanted into the hormonally synchronized recipient cows or ewes. For chimeric blastocyst, ten haSCs labelled by EGFP fluorescence were microinjected into an 8-cell stage embryo. The reconstructed embryos were cultured in FACE medium for 4-5 h, then changed into the G1/G2 medium (1:1; Vitrolife) to obtain the chimeric blastocysts.

### Embryoid body formation assay

Embryoid body (EB) formation was performed as previously described with slight modification ^48^. Briefly, haSCs were dissociated into single cells with TrypLE (Thermo) and cultured for 7 days on ultra-low attachment plates in Iscove’s modified Dulbecco’s medium (IMDM; Thermo) supplemented with 1% nonessential amino acids (Thermo), 1% penicillin-streptomycin (Thermo), and 15% FBS (Gibco). Subsequently, EBs were collected and dissociated by 0.05% trypsin-EDTA (Thermo) at 37 °C for 10 min. After enrichment, cells were plated on matrigel-coated slides for 3 days before IF staining.

### Teratoma formation assay

Teratoma formation was performed as previously described ^34^. In brief, nearly 10^9^ haSCs were subcutaneously injected into the hind limbs of 8-week-old nude mice. After 5 weeks, fully formed teratomas were dissected and fixed with PBS containing 4% PFA, then embedded in paraffin, sectioned, and stained with hematoxylin and eosin (H&E) for histological analysis.

### Intracytoplasmic haSCs injection

Angus cattle and Dorper sheep ovaries were obtained from commercial slaughterhouses, and transported to the laboratory within 2 h at 28 ℃ in sterile saline solution (8.5 mg/mL NaCl) with a thermal container. The oocytes, which possessed at least three layers of cumulus, were selected for IVM according to the methods described above. Intracytoplasmic haSCs injection (iCHI) was performed following the published protocols with slight modifications ^1, 2, 48^. Briefly, haSCs were arrested metaphase by culturing in FACE medium containing 0.05 μg/mL demecolcine for 12 h. Following centrifugation, cells were rinsed three times with PBS for removal of the demecolcine, and resuspended in the M2 medium containing 10 μM Y-27632 (Stemcell) and 1.5% (w/v) polyvinylpyrrolidone (PVP). Each nucleus from haSCs was directly injected into an IVM oocyte using the Piezo micromanipulator (Primetech) in the M2 medium containing 5 μg/mL cytochalasin B (CB) on a 37 °C heating stage of an inverted microscope (Nikon). The reconstructed embryos were cultured in KSOM-AA medium (Millipore) for 1 hr and then activated using the previously described protocols with slight modifications ^13, 30, 44, 52^. Briefly, the reconstructed embryos were treated with 5 μM ionomycin in Ca^2+^-free KSOM-AA medium for 5 min, followed by incubation in KSOM-AA medium containing 10 μg/mL cycloheximide (CHX) for 5 h. After activation, the embryos were rinsed several times in M2 medium and cultured in KSOM-AA medium supplemented with 3 mg/mL of BSA under mineral oil at 38.5 °C, 5% CO_2_, 5% O_2_, and 90% N_2_ for 3 days. Then, the embryos were transferred into KSOM-AA medium containing 5% FBS (Gibco) for further cultivation.

### Embryo Transfer

For bovine embryo transfer, blastocysts were stored in the TCM-199 medium (Thermo) supplemented with 15% FBS (Gibco). The iCHI blastocysts were transferred via the non-surgical transcervical method as previously described ^53^. Briefly, a single blastocyst was transferred into the uterine horn of hormonally synchronised (CIDR progesterone + cloprostenol + gonadorelin) nulliparous heifer at day 7 (estrus day 0 = day of micromanipulation). For ovine embryo transfer, 2-cell stage embryos were stored in TCM-199 medium (Thermo) supplemented with 10% BSA. The ovine embryos were transferred via surgical method using a TomCat catheter (Sovereign). Eight iCHI embryos were transferred into the oviduct of hormonally synchronized (CIDR progesterone + FSH + cloprostenol) recipient ewes at day 2 (estrus day 0 = day of micromanipulation). Pregnancy detection was diagnosed on day 35 and 50 of bovine and ovine embryonic development, respectively, using transrectal ultrasound EVO scanner (5.0MHz linear probe, E.I. Medical Imaging).

### Reverse transcription-quantitative PCR

Reverse transcription-qPCR assays were performed according to the Minimum Information for Publication of Quantitative Real-Time PCR Experiments (MIQE) guidelines ^54^. Briefly, total RNA was extracted from cells by a single-step method using FreeZol Reagent (Vazyme) according to the manufacturer’s protocol. The concentration and purity of RNA were determined using Nanodrop spectrophotometry (Thermo) to ensure the 260/280 ratio was within 1.8-2.0 range. The RNA integrity was further verified by denaturing agarose gel electrophoresis. Genomic DNA was removed using gDNA wiper Mix, followed by cDNA synthesis using HiScript II Q RT SuperMix for qPCR (+gDNA wiper; Vazyme) according to the manufacturer’s protocol. The real-time quantitative PCR reaction was performed using ChamQ Universal SYBR qPCR Master Mix Kit (Vazyme), signals were detected with Applied Biosystems QuantStudio 3 (Thermo) analyzed by the QuantStudio design & analysis software (v1.5.1) following the manufacturers’ instruction manual. Thermocycling conditions were 95 °C for 30 sec, followed by 40 cycles of 95 °C for 10 sec and 60 °C for 30 sec. No template controls (NTCs) were performed using the carrier-containing reaction mix and no template DNA, in all cases no amplification occurred. Results were generated from 3 technical and 3 biological replicates for each condition. Relative gene expression analysis was normalized to internal glyceraldehyde-3-phosphate dehydrogenase (GAPDH) or beta-actin gene, and measured using the 2^(-ΔΔCt)^ method. The primer information was presented in Supplementary Table 6; The reaction efficiencies, validations, and MIQE checklist were presented in Supplementary Fig. 7 and Table 8.

### Bisulphite sequencing

Genomic DNA was extracted using the tissue and blood DNA extraction kit (Qiagen) and treated with the Methylamp DNA modification kit (EpiGentek). To obtain sperm genomic DNA, samples were pretreated with dithiothreitol (DTT) for 3 hr, following proteinase K lysis and phenol-chloroform extraction. The bisulfite conversion was performed as described previously ^55^. The PCR products were cloned into vectors using TA/Blunt-Zero Cloning Kit (Vazyme), and individual clones were sequenced. The primer information is presented in Supplementary Table 6.

### Plasmid construction

For the PRDM1-tdTomato (PT) and TFAP2C-mNeonGreen (TG) reporter plasmid, construction was performed according to the recently reported protocol ^51^. Oligonucleotides encoding single-guide RNAs (sgRNAs) were inserted into BbsI-digested pX330 plasmid (Addgene #42230) to create CRISPR/Cas9 expression vectors. For the episomal PE plasmid, the coding sequence of Cas9 (H840A)::M-MLVrt (D200N, T306K, W313F, T330P, L603W), derived from Prime editor 2 system (PE2, Addgene #132775) ^19^, was codon-optimized based on the general codon usage in cattle and sheep then synthesized by commercial company. The above vector was designated boPE2 and ovPE2. Indeed, “all-in-one” episomal PE plasmid is the same as epiCRISPR ^56^ except that the Cas9 was replaced by boPE2 or ovPE2. All CRISPR editor plasmids used in this work were assembled using the USER cloning method as previously described ^19^. DNA amplification was conducted by Phanta Max Super-Fidelity DNA Polymerase (Vazyme). Integrate the synthesised DNA into the designated positions on the vector using ClonExpress Ultra One Step Cloning Kit V2 (Vazyme). The pegRNA targeting gene was designed by the online program (http://pegfinder.sidichenlab.org) ^57^ based on the GenBank sequence. For the episomal doxycycline (dox)-inducible PRM1-EGFP plasmid, construction was performed as previously described with slight modification ^17, 58^. Briefly, the sequence of mouse PRM1 (GenBank NP_038665.1), oriP-EBNA1 (Addgene #84031), and EGFP (Addgene #176015) was synthesized by commercial company, and inserted into pCW57-MCS1-P2A-MCS2 plasmid (Addgene #89180). See more details in Supplementary Fig. 8.

### Generation of BT::TG reporter b-haSCs

The culture dishes were prepared using the methods described above, but the multidrug-resistant DR4 MEFs (Gibco) were used as the feeder cells. The transgenic manipulation was performed as previously described with slight modification ^51^. First, to knock in the TFAP2C-mNeonGreen (TG) reporter, approximately 10^4^ b-haSCs were disassociated using TrypLE (Thermo) and transfected with plasmids. A total of 2 μg CRISPR/Cas9 plasmids and TG reporter plasmids were transfected into b-haSCs using Lipomaster 3000 Transfection Reagent (Vazyme) following the manufacturer’s protocol. After transfection 72 h, b-haSCs were selected by 0.2 μg/mL puromycin. The colonies of puromycin-resistant b-haSCs were picked, and correct targeting was determined by PCR genotyping. Similarly, the PRDM1-tdTomato (PT) reporter was further knocking into the TG b-haSCs. After selection with 400 μg/mL G418, colonies were picked and correct targeting was determined by PCR genotyping. The gRNAs and primers information were presented in Supplementary Table 6.

### Generation of MSTN-edited haSCs

The culture dishes were prepared using the methods described above, but the multidrug-resistant DR4 MEFs (Gibco) were used as the feeder cells. Perform a 100% medium change daily and passage cells every 5 days. Genome editing was performed as previously described with slight modification ^3, 56^. In brief, approximately 10^4^ haSCs were disassociated using TrypLE (Thermo) and transfected with plasmids. A total of 2 μg ePE plasmid were transfected into haSCs using Lipomaster 3000 Transfection Reagent (Vazyme) following the manufacturer’s protocol. After transfection 72 h, haSCs were selected by 0.2 μg/mL puromycin. For analysis of single cell-derived clones, the haSCs were disaggregated into single cells at day 15 post-transfection, and 1,000 cells were seeded on one well of the 6-well dishes with puromycin-free FACE medium for sub-colony formation. After 14 days, singular sub-colonies were picked and evaluated by PCR genotyping. Editing sites of MSTN gene according to previous reports ^59, 60^. The primers and pegRNA information is presented in Supplementary Table 6, 7.

### Genotyping genome-edited mutations

Genotyping was performed as previously described with slight modifications ^56, 61^. Briefly, the target sites were amplified by PCR with specific primers from genomic DNA using Phanta Max Super-Fidelity DNA Polymerase (Vazyme). The amplicon was cloned into the pCE3 plasmid using the Ultra-Universal TOPO Cloning Kit (Vazyme) and transformed to competent *E. coli* strain DH5α. After overnight culture at 37 °C, fifteen emerged colonies were randomly picked out, and underwent Sanger sequencing at Sangon Biotech Co., Ltd., and the results were analyzed using TIDE software ^62^. The percentage of edited clones was calculated as the number of clones detected to possess at least one nucleotide change relative to the total number of sequenced clones. Mutations were identified by aligning the sequenced amplicon to the wild-type target sequence. The primer information is presented in Supplementary Table 6.

### WGS and data processing

Whole genome sequencing (WGS) was performed as described previously ^48,63^. Briefly, libraries were generated from 1 mg of genomic DNA using the TruSeq DNA Sample Preparation Kit (Illumina), and sequenced on the HiSeq 4000 system (Illumina) by Lc-Bio Technologies Co., Ltd. For data processing, copy number variation (CNV) detection was performed as previously described ^64^. The template genome was downloaded from NCBI RefSeq, including cattle (ARS-UCD2.0) and sheep genome (ARS-UI_Ramb_v3.0).

### WGBS and data processing

Whole genome bisulfite sequencing (WGBS) was performed according to previously ^48, 65^. Briefly, genomic DNA was extracted from haSCs using the DNeasy Blood & Tissue extraction kit (Qiagen) following end repair and A-tailing. The DNA was fragmented into around 200-300 bp by ultrasonicator (Covaris). Bisulfite-converted DNA was captured using EZ-96 DNA Methylation-DirectTM MagPrep (Zymo) according to the manufacturer’s instructions. Library sequencing was performed on the HiSeq 4000 system (Illumina) by Lc-Bio Technologies Co., Ltd. The cleaned data were aligned to the cattle (ARS-UCD2.0) reference with Bismark/Bowtie2 mode. Bismark2summary command was used to calculate the genome CpG methylation level. RnBeads R package was used to calculate the percentage of methylation.

### CUT&Tag and data processing

CUT&Tag was performed using the Hyperactive Universal CUT&Tag Assay Kit for Illumina (Vazyme) according to the manufacturer’s instructions. Paired-end sequencing was further performed on the HiSeq 4000 system (Illumina) by Frasergen Co., Ltd. The data processing was performed as described previously ^66^. Briefly, paired-end reads were aligned using Bowtie2 (2.2.5) with options: -local-very-sensitive-local–no-unal–no-mixed–no-discordant-phred33 -I 10 -X 700. For peak calling, the parameters used were MACS2 callpeak -t input_file -p 1e-5 -f BEDPE/BED -keep-dup all -n out_name. The antibody information is presented in Supplementary Table 5.

### buRNA-seq and data processing

The bulk RNA sequencing (buRNA-seq) was followed by previously published studies ^34, 49^. Briefly, total RNA was extracted from haSCs using TRIzol Plus RNA Purification Kit (Thermo) following the manufacturer’s instructions, and the RNA integrity was checked using Bioanalyzer 2100 (Agilent). The total cDNA library was then amplified by 18-20 cycles for library construction following the manufacturer’s instructions (Illumina, USA). Sequencing was further performed on the HiSeq 4000 system (Illumina) by Frasergen Co., Ltd. After removing low-quality reads and adapters, the raw reads were mapped to the cattle (ARS-UCD2.0) or sheep (ARS-UI_Ramb_v3.0) genome using Tophat (v1.3.3) with default parameters ^67^. Expression levels for all RefSeq transcripts were quantified to fragments per kilobase of exon model per million mapped reads (FPKM) using Cufflinks (v1.2.0) ^68^. Gene Ontology (GO) analyses were performed using R package GO.db and Database for Annotation, Visualization, and Integrated Discovery (DAVID) ^69^.

### scRNA-seq and data processing

The single-cell RNA sequencing (scRNA-seq) was followed by previously published studies ^65, 70^. Briefly, haSCs or E8∼E14 embryos were dissociated into single cells with TrypLE (Thermo) and resuspended in 1 Ca^2+^-Mg^+^ free PBS containing 0.04% BSA. Next, 300-1000 living cells were loaded on a Chromium Single Cell Controller (10x Genomics) to generate single-cell gel beads in emulsion (GEMs) cDNAs. Then, recovered cDNAs were used to construct a library using Single Cell 3’ Library and Gel Bead Kit V2 (10x Genomics) according to the manufacturer’s instructions. Library sequencing was performed on the HiSeq 4000 system (Illumina) with pair-end 150 bp (PE150) by Lc-Bio Technologies Co., Ltd. The raw sequencing data were aligned and quantified using the Cell Ranger (10x Genomics) pipeline against the cattle (ARS-UCD2.0) reference genome. The Seurat R package was used to merge the scRNA-seq and cell clustering. Gene expression was calculated using the LogNormalize method in the NormalizeData function. RunPCA function was used in principal component analysis (PCA) with the most variable genes. Cell clusters were identified by a shared nearest neighbor (SNN) modularity optimisation based on a clustering algorithm with the FindClusters function. The *t*-SNE (*t*-distributed stochastic neighbor embedding) dimensionality reduction was generated using the RunTSNE function and the UMAP (uniform manifold approximation and projection) was performed using the RunUMAP function. Differentially expressed genes were identified using FindAllMarkers, and the top 30 differentially expressed genes (DEGs) were marked as signature genes. Heatmaps were plotted by the DoHeatmap function.

## Data availability

The high-throughput sequencing data generated during this study were deposited to the National Center for Biotechnology Information (NCBI) Gene Expression Omnibus (GEO) database under the SuperSeries accession GSE250497. Other data that support the findings of this study are available within the article and its Supplementary Information, Source data, or from the corresponding author upon reasonable request.

## Acknowledgements

We thank Hong Wang (Inner Mongolia University) for technical help. This study was supported by the National Natural Science Foundation of China (32341052, 32360837), Scientific and Technological Innovation 2030 (2023ZD0404803), Inner Mongolia Open Competition Projects (2022JBGS0025), Inner Mongolia Science and Technology Leading Team (2022LJRC0006), Inner Mongolia Science and Technology Major Projects (2021ZD0009, 2021ZD0008, 2022ZD0008, 2023KJHZ0028), Inner Mongolia Young Talents Projects (NJYT23138), Inner Mongolia Natural Science Foundation (2023MS03004), Central Government Guides Development (2022ZY0212), National Agricultural Science and Technology Project (NK2022130203), Collaborative Innovation among Universities in Hohhot (XTCX202306), Ministry of Education Engineering Centre Project (JYBGCSYS2022), Xinjiang Uygur Science and Technology Major Project (2023A0201116), TongLiao Open Competition Projects (TL2024TW0020103).

## Author contributions

L.Y., A.D., X.L., L.S., D.W., S.W., Z.H., L.B., C.B., G.S., Z.W., L.Z. performed experiments. L.Y., G.L., and S.G. designed experiments, L.Y. and G.L. wrote the manuscript.

## Additional information

**Correspondence and requests for materials** should be addressed to Lei Yang, Shaorong Gao, or Guangpeng Li.

