## Extended Data for "One-step generation of modified cattle and sheep from spermatid-like haploid stem cells"

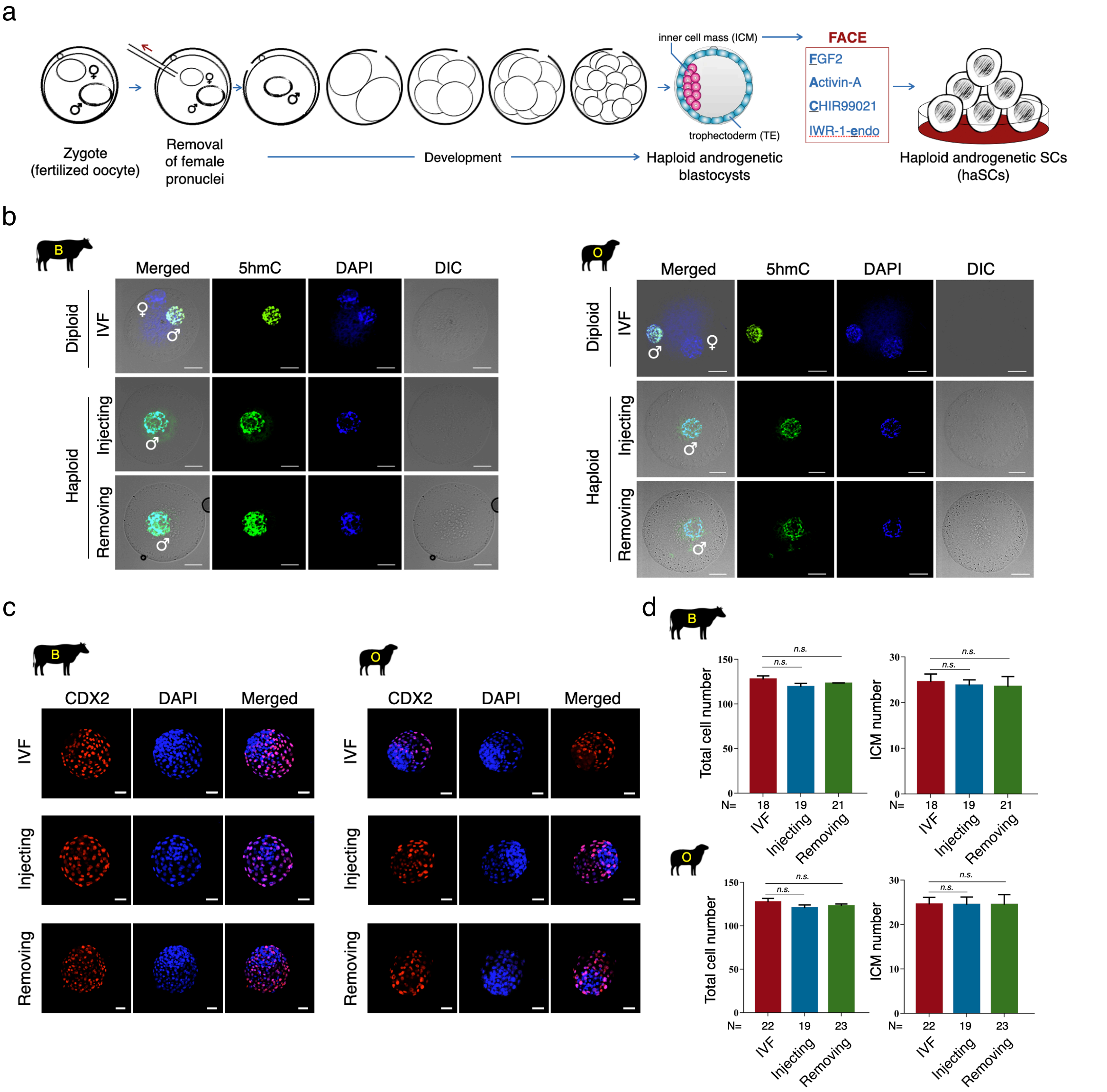


### Extended Data Fig. 1

**Generation of ruminant haploid androgenetic embryos.**

a. Schematic overview of haSCs derivation, haploid embryos were generated by removing female pronucleus from normal diploid zygotes. ♂, male pronucleus; ♀, female pronucleus.

b. Immunostaining for 5-hydroxymethylcytosine (5hmC) in bovine (left) and ovine (right) embryos. The haploid androgenetic embryos are generated by sperm injection into enucleated oocytes or by removing female pronucleus from fertilized oocytes. The diploid IVF embryos were used as controls. ♂, male pronucleus; ♀, female pronucleus; Scale bar, 25 µm.

c. Immunostaining for trophectoderm marker CDX2 in the bovine and ovine blastocysts on day 7 after activation or insemination. The haploid and diploid embryos are generated as described in Extended Data Fig. 1b. Scale bar, 50 µm.

d. Quantification for the cell numbers in the bovine and ovine blastocysts on day 7 after activation or insemination. The haploid and diploid embryos are generated as described in Extended Data Fig. 1b; ICM, inner cell mass; N, the total number of blastocysts analyzed for each group; mean ± s.d., n = 3 independent experiments; n.s., not significant by Student's *t*-test.

a

| | | 2i/L | 4i | t2iL+Gö | 5i/L/A | ABCL | LCDM | EPSC | CTFR | bEPSCM |
|---|---|---|---|---|---|---|---|---|---|---|
| Inhibitors | GSK3 | CHIR99021 (C) | CHIR99021 (C) | CHIR99021 (C) | IM-12 (I) | CHIR99021 (C) | CHIR99021 (C) | CHIR99021 (C) | | CHIR99021 (C) |
| | MEK | PD0325901 (P) | PD0325901 (P) | PD0325901 (P) | PD0325901 (P) | | | PD0325901 (P) | | |
| | P38 | | SB203580 (580) | | | | | SB203580 (580) | | |
| | JNK | | SP600125 (125) | | | | | TCSJNK60 (T) | | |
| | PKC | | | Gö6983 (G) | | | | | | |
| | WNT | | | | | | | XAV939 (X) | IWR-1-endo (E) | XAV939 (X) or IWR-1-endo (E) |
| | ROCK | | Y27632 (Y) | | Y27632 (Y) | | | | | |
| | BRAF | | | | SB590885 (885) | | | | | |
| | SRC | | | | WH-4-023 (W) | | | A-419259 (259) | | WH-4-023 (W) |
| | Histamine | | | | | | Dimethindene maleate (D) | | | |
| | PARP1 | | | | | | Minocycline hydrochloride (M) | | | |
| Growth factors | LIF (L) | ✓ | ✓ | ✓ | ✓ | ✓ | ✓ | ✓ | | ✓ |
| | FGF2 (F) | | ✓ | | | | | | ✓ | ✓ |
| | TGFβ (T) | | ✓ | | | | | | | ✓ |
| | Activin-A (A) | | | | ✓ | ✓ | | | | ✓ |
| | BMP4 (B) | | | | | ✓ | | | | |
| | Vitamin C (V) | | | | | | | | | ✓ |
| | Original application | M | | | | | M | M P | B O | B |
| | Reference PMID | 18497825 | 25543152 | 25215486 | 25090446 | 29076502 | 28388409 | 29019987 31160711 | 29440377 33065542 | 33833056 |

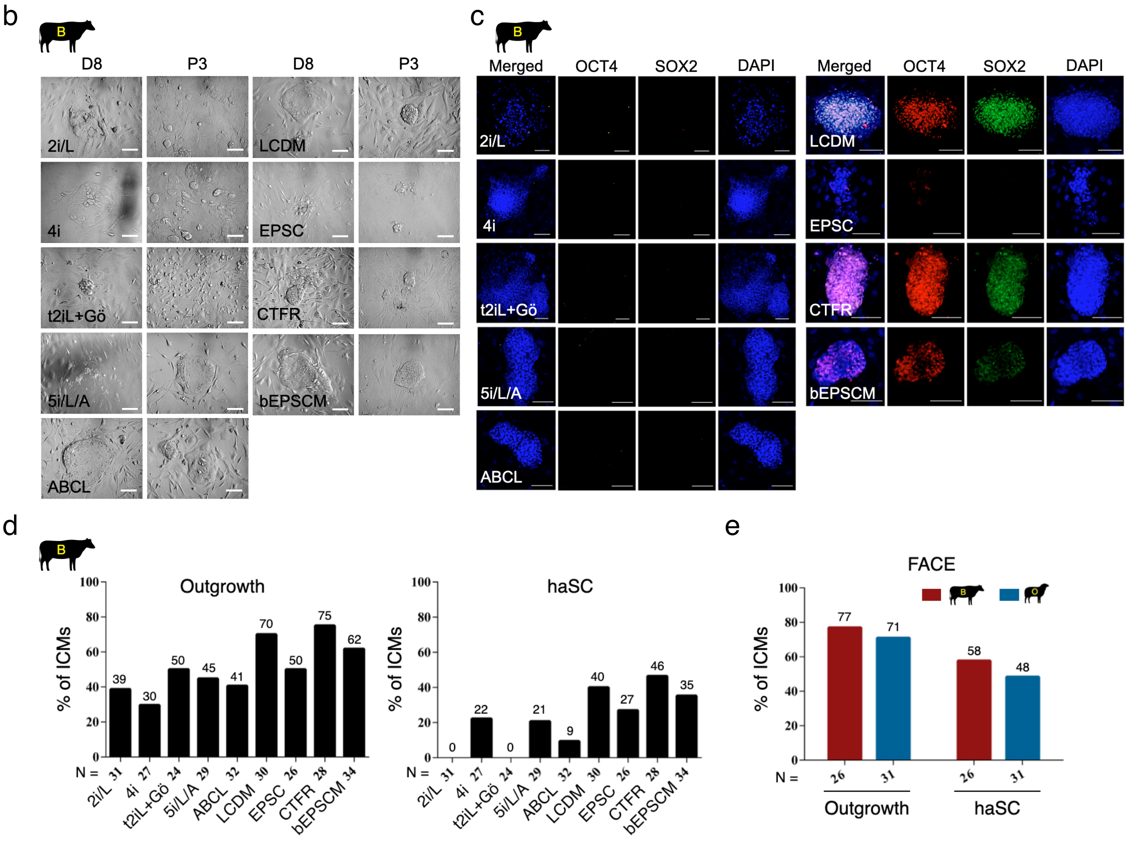


### Extended Data Fig. 2

**Derivation of ruminant haSCs from haploid androgenetic embryos.**

a. List of main components in the well-known medium that support the derivation of mouse, human, and bovine diploid SCs, including 2i/LIF, mTeSR1, t2iL+Gö, 5i/L/A, ABCL, LCDM,

EPSC, CTFR, and bEPSCM. The brief names of these chemicals and their respective pathways which they regulate are also shown.

b. Morphology of bovine haploid androgenetic ICMs under different culture medium conditions at indicated time points. D, day; P, passage; Scale bars, 200 μm.

c. Representative immunofluorescence (IF) images of pluripotency factors in the indicated primary outgrowth. Scale bar, 100 μm.

d, e. The efficiency of outgrowth and haSCs derivation of haploid androgenetic ICMs by different culture mediums. N = total number of ICMs used for each condition.

a

| No. | Small molecucules | No. | Small molecucules | No. | Small molecucules | No. | Small molecucules | No. | Small molecucules |
|---|---|---|---|---|---|---|---|---|---|
| 1 | FCLE | 6 | FCWE | 11 | FALD | 16 | FAXD | 21 | FACD |
| 2 | FDLE | 7 | FCWL | 12 | FALM | 17 | FACL | 22 | FACV |
| 3 | FWLE | 8 | FCDL | 13 | FAWT | 18 | FACT | 23 | FACM |
| 4 | FMLE | 9 | FCML | 14 | FAET | 19 | FACX | 24 | CEWL |
| 5 | FMWE | 10 | FCXL | 15 | FAXT | 20 | FACE | 25 | CTML |

CHIR99021 (C); Dimethindene maleate (D); Minocycline hydrochloride (M); LIF (L); IWR-1-endo (E); FGF2 (F); XAV939 (X); WH-4-023 (W); TGFβ (T); Activin-A (A); Vitamin C (V)

b B

| Combi | No. of embryos | Outgrowth rat | Cell lines |
|---|---|---|---|
| 1 | 25 | 2/25 | 1/2 |
| 2 | 21 | 1/21 | 0 |
| 3 | 24 | 3/24 | 0 |
| 4 | 23 | 2/23 | 0 |
| 5 | 27 | 1/27 | 0 |
| 6 | 24 | 4/24 | 0 |
| 7 | 22 | 2/22 | 0 |
| 8 | 19 | 2/19 | 0 |
| 9 | 17 | 1/17 | 0 |
| 10 | 21 | 1/21 | 0 |
| 11 | 22 | 7/22 | 0 |
| 12 | 27 | 6/27 | 0 |

| Combi | No. of embryos | Outgrowth rat | Cell lines |
|---|---|---|---|
| 13 | 16 | 3/16 | 0 |
| 14 | 14 | 4/14 | 0 |
| 15 | 18 | 6/18 | 0 |
| 16 | 23 | 9/23 | 1/9 |
| 17 | 17 | 4/17 | 0 |
| 18 | 12 | 10/12 | 7/10 |
| 19 | 24 | 21/24 | 19/21 |
| 20 | 27 | 25/27 | 24/25 |
| 21 | 21 | 15/21 | 1/15 |
| 22 | 17 | 6/17 | 0 |
| 23 | 16 | 5/16 | 0 |
| 24 | 19 | 0/19 | 0 |
| 25 | 15 | 1/15 | 0 |

c

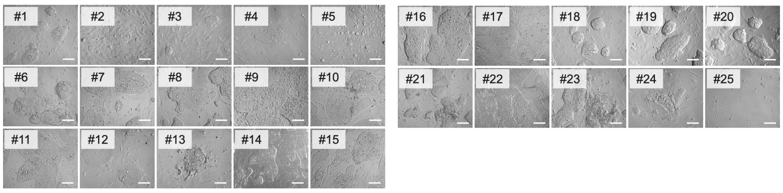


d

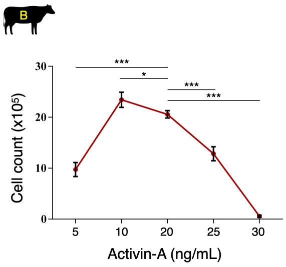


e

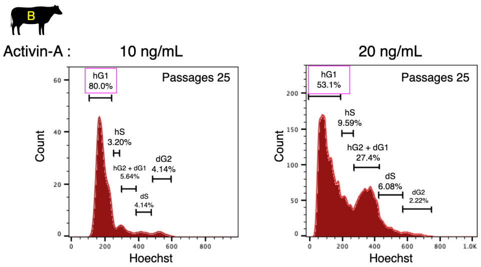


**Extended Data Fig. 3**

**FACE medium supports ruminant haSCs long-term culture.**

a. List of different combinations of small molecules used in this study.

b. Summary of b-haSCs derivation efficiency from bovine haploid androgenetic ICMs by different combinations of small molecules.

c. Morphology of bovine haploid androgenetic ICMs under different combinations of small molecules. Scale bars, 200 μm.

d. The total cell numbers of b-haSCs during cell passaging under different concentrations of

Activin-A. Cells were plated at $5 \times 10^5$ cells per well and were cultured for 4 days; mean ± s.d.; n = 3 independent experiments; **P* < 0.05, ****P* < 0.001 as compared with 20 ng/mL group, by two-tailed Student's *t*-test.

e. Haploidy analysis in b-haSCs derived using 10 ng/mL and 20 ng/mL of Activin-A. The figure on the left is the same as Fig. 1e; n = 3 independent experiments with similar results. Note that 10 ng/mL Activin-A was beneficial for maintaining haploidy stability.

a

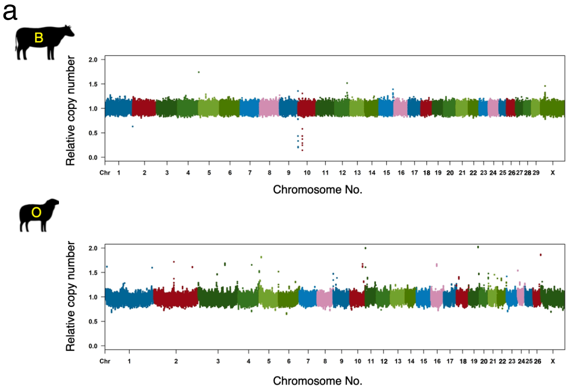


b

| Species | Passage number | Normal haploid karyotype / Total (%) |
|---|---|---|
| B | 25 | 31/31 (100.0) |
| | 28 | 33/33 (100.0) |
| | 31 | 32/32 (100.0) |
| O | 25 | 30/30 (100.0) |
| | 28 | 31/31 (100.0) |
| | 31 | 29/30 (96.7) |

c

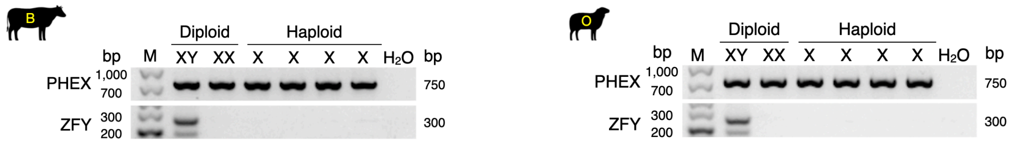


**Extended Data Fig. 4**

**Characterization of ruminant haSCs.**

a. DNA copy number variation (CNV) analysis of ruminant haSCs at passage 25. Note that no significant genomic alternations. The results were displayed on a log2 scale. Chromosomes are arranged in numerical order and in different colours.

b. Summary of karyotyping analysis of ruminant haSCs at different passages. Total, the total number of examined cells in which chromosome was successfully spread.

c. Determination of the sex chromosome of ruminant haSCs by PCR assay. The primers are specific for X chromosome-specific PHEX and Y chromosome-specific ZFY genes. Normal XX and XY diploid SCs were used as controls. M, DNA marker.

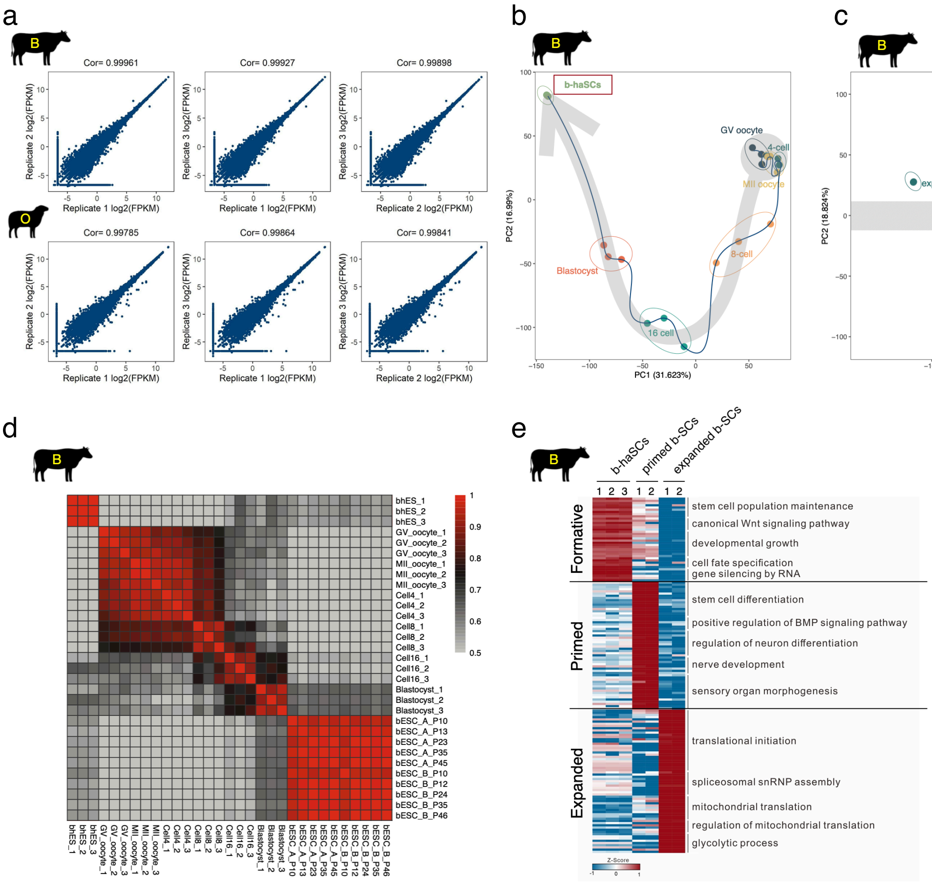


**Extended Data Fig. 5**

**Transcriptome analysis of ruminant haSCs.**

a. Scatter-plots showing the reproducibility of buRNA-seq between different biological replicates of ruminant haSCs. The Pearson's correlation coefficients (Cor.) are shown.

b. Principal-component analysis (PCA) showing the separation of b-haSCs and bovine pre-implantation embryos.

c. Scatter plot based on PCA of b-haSCs, diploid primed b-SCs, and diploid expanded b-SCs, showing that the projection position of b-haSCs is located between expanded and primed b-SCs.

d. Correlation matrices showing coefficients among b-haSCs, diploid primed b-SCs, diploid expanded b-SCs, and pre-implantation bovine embryos.

e. Heatmap displaying the differentially expressed genes (DEGs) among b-haSCs, primed b-SCs, and expanded b-SCs ($P < 0.05$, 2-fold change; left). Gene Ontology (GO) terms about the biological processes enriched in DEGs are listed (right).

RNA-seq data of b-haSCs generated in this study, previously published data for bovine diploid pre-implantation embryos are from Graf A., *et al.*, Proc Natl Acad Sci U S A. 2014, PMID: 24591639 (GSE52415), diploid primed b-SCs are from Bogliotti Y., *et al.*, Proc Natl Acad Sci U S A. 2018, PMID: 29440377 (GSE110040), diploid expanded b-SCs are from Zhao L., *et al.*, Proc Natl Acad Sci U S A. 2021, PMID: 33833056 (GSE129760).

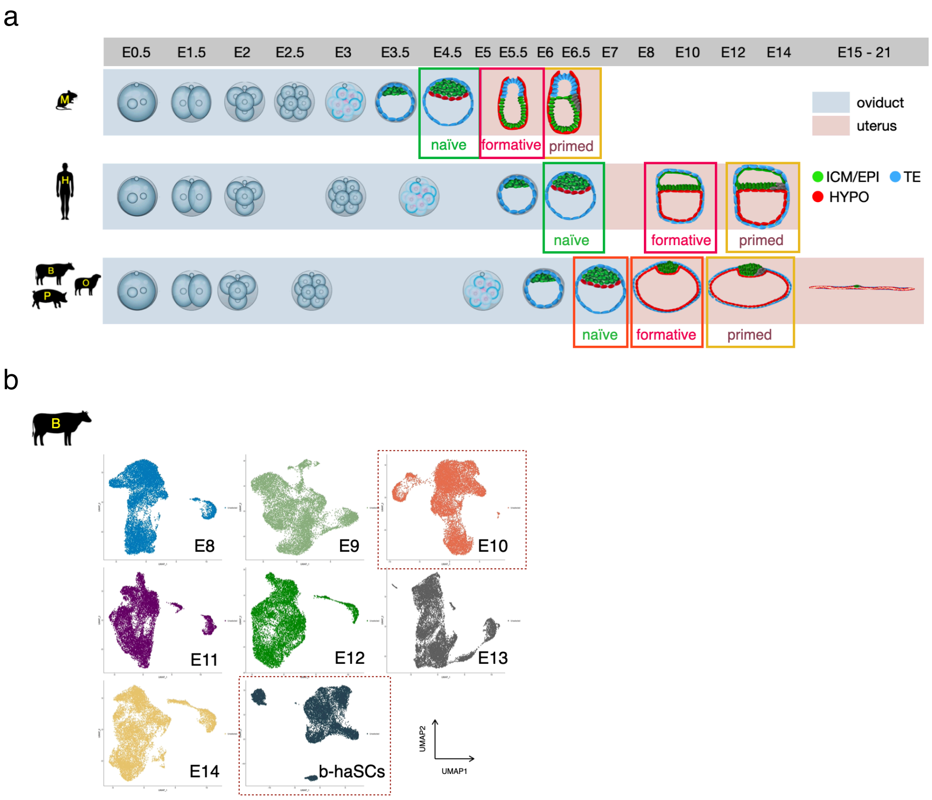


**Extended Data Fig. 6**

**Ruminant haSCs exhibit transcriptional features of formative pluripotency.**

a. Developmental progression of pluripotency in embryos during peri-implantation. Formative pluripotency may be present at the early stages of gastrulation, according to developmental time and lineage segregation of mouse, monkey, pig, and human embryos (Smith A., Development. 2017, PMID: 28143843; Peng G., *et al.*, Dev Cell. 2016, PMID: 27003939; Zhi M., *et al.*, Cell Res. 2022, PMID: 34848870; Wen J., *et al.*, J Biol Chem. 2017, PMID: 28298438; Nakamura T., *et al.*, Nature. 2016, PMID: 27556940; van Leeuwen J., *et al.*, PLoS One. 2015, PMID: 26076128; Pérez-Gómez A., *et al.*, Front Vet Sci. 2021, PMID: 34212020.). E, embryonic day; ICM, inner cell mass; EPI, epiblast; HYPO, hypoblast; TE, trophectoderm.

b. A uniform manifold approximation and projection representation (UMAP) of bovine peri-implantation embryos (E8~E14) and b-haSCs sequenced by 10x Genomics.

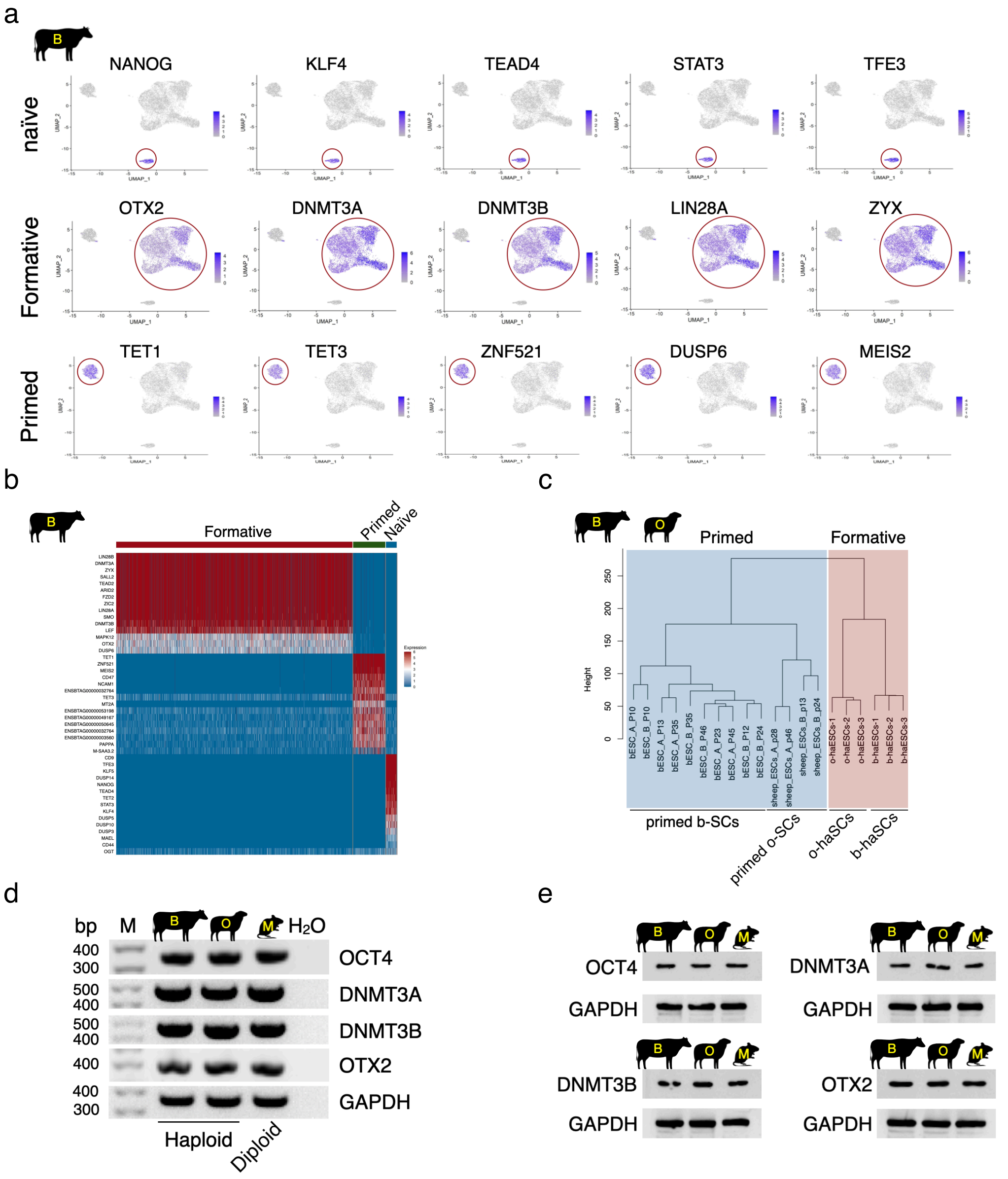


### Extended Data Fig. 7

**Further analyses of ruminant haSCs about formative features.**

a. The expression of typical naïve-, formative-, and primed-pluripotency genes in b-haSCs at the single-cell level.

b. Heatmap displaying the expression of naïve-, formative-, and primed-pluripotency genes in b-haSCs sequenced by scRNA-seq.

c. Hierarchical clustering analysis of b-haSCs, o-haSCs, primed b-SCs, and primed o-SCs based on the global transcriptome.

d. RT-PCR analysis of core formative markers (DNMT3A, DNMT3B, OTX2) and pluripotency marker (OCT4) in b-haSCs, o-haSCs, and formative m-SCs. M, DNA marker.

e. Western blots analysis of core formative markers (DNMT3A, DNMT3B, OTX2) and pluripotency marker (OCT4) in b-haSCs, o-haSCs, and formative m-SCs.

RNA-seq data of o-haSCs and b-haSCs generated in this study, previously published data for bovine primed-b-SCs are from Bogliotti Y., *et al.*, Proc Natl Acad Sci U S A. 2018, PMID: 29440377 (GSE110040), diploid primed o-SCs are from Vilarino M., *et al.*, Reproduction. 2020, PMID: 33065542 (PRJNA609175).

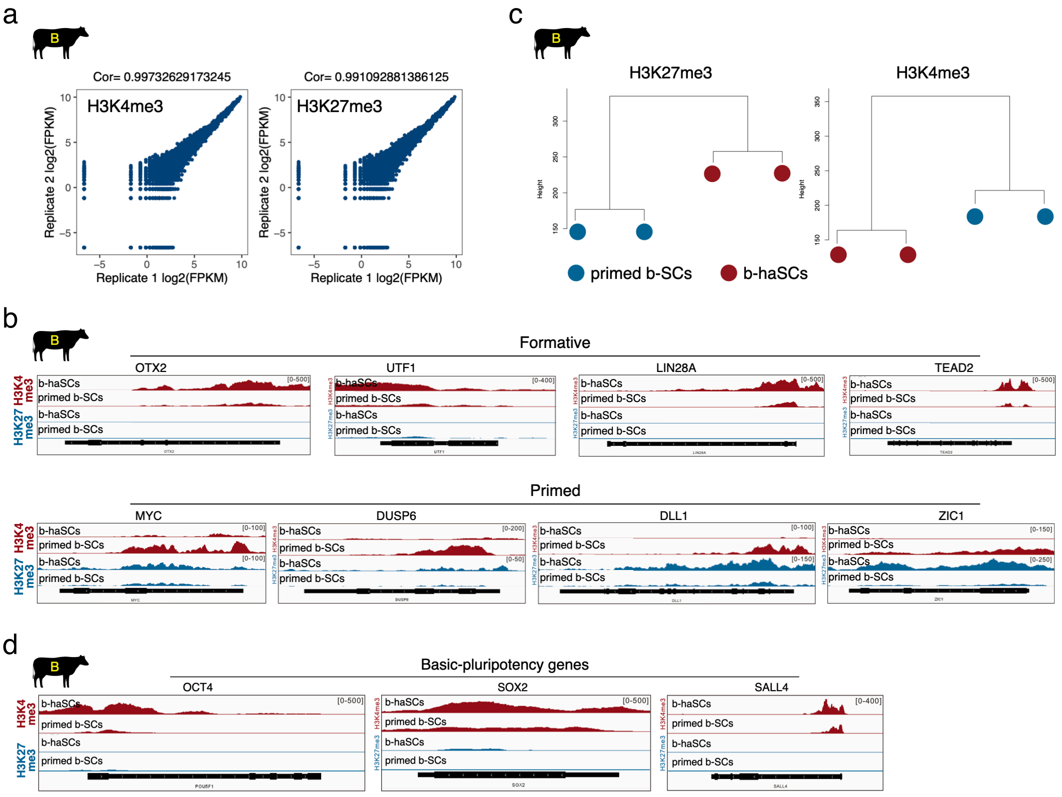


### Extended Data Fig. 8

**Ruminant haSCs display epigenetic signature of formative pluripotency.**

a. Scatter-plots showing the reproducibility of CUT&Tag assay between different biological replicates of b-haSCs. The Pearson's correlation coefficients (Cor.) are shown.

b. UCSC genome browser views showing H3K4me3 and H3K27me3 tracks of representative formative- and primed-pluripotency genes in b-haSCs and primed b-SCs.

c. Hierarchical clustering analysis of b-haSCs and primed b-SCs based on the H3K4me3 and H3K27me3 CUT&Tag signals.

d. UCSC genome browser views showing H3K4me3 and H3K27me3 tracks of selected pluripotency-associated genes in b-haSCs and primed b-SCs. Note that H3K4me3 but not H3K27me3 signals exhibited high enrichment levels in both b-haSCs and primed b-SCs.

The H3K4me3 and H3K27me3 data of b-haSCs generated in this study, previously published data for diploid primed b-SCs are from Bogliotti Y., *et al.*, Proc Natl Acad Sci U S A. 2018, PMID: 29440377 (GSE110040).

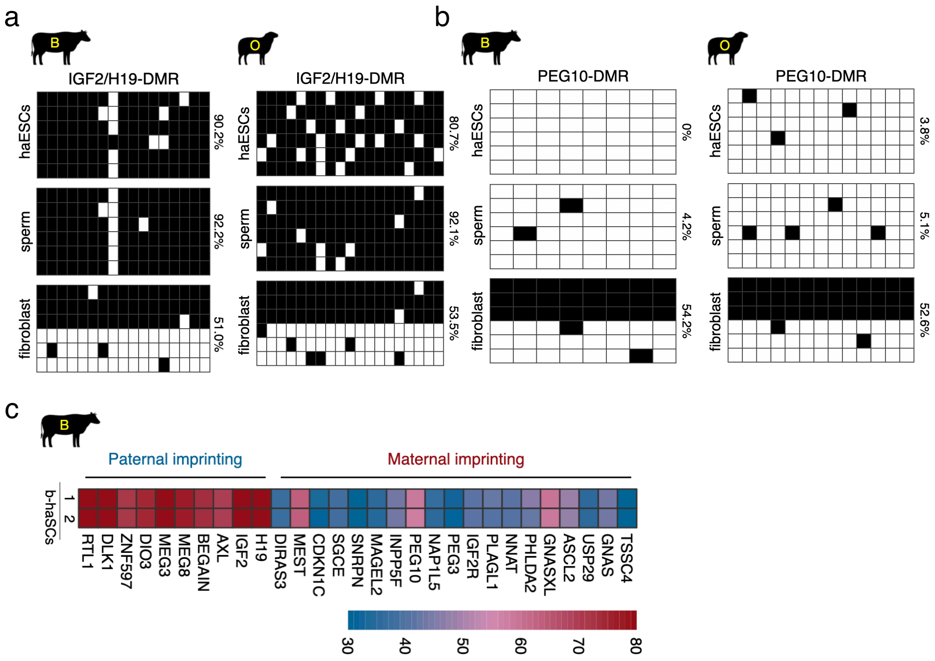


**Extended Data Fig. 9**

**Ruminant haSCs retain parental imprinting states.**

a, b. Bisulfite pyrosequencing analysis of paternally (a) and maternally (b) imprinted regions in b-haSCs and o-haSCs. The filled and open squares represent methylated and unmethylated CpG sites, respectively.

c. Whole genome bisulfite sequencing (WGBS) analysis of known imprinting control regions in b-haSCs.

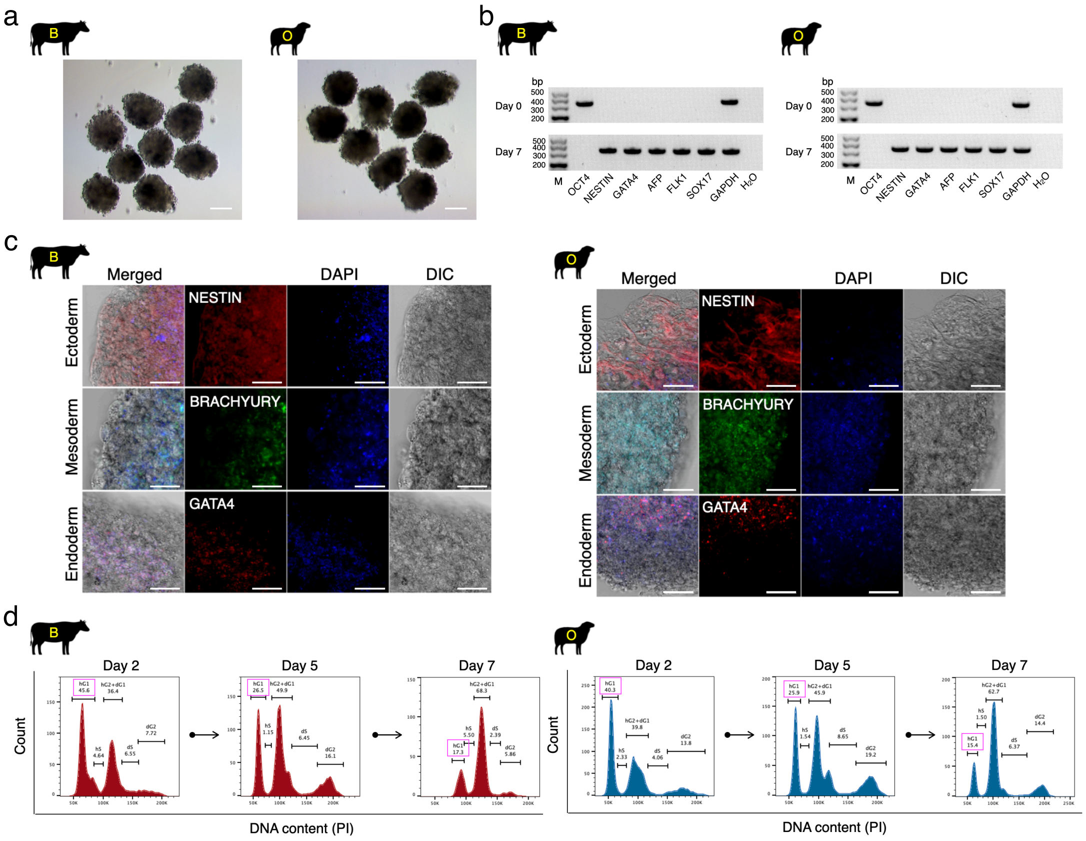


### Extended Data Fig. 10

**Ruminant haSCs harbour potency for the formation of embryoid body *in vitro*.**

a. Morphology of embryoid bodies (EBs) derived from ruminant haSCs. Scale bar, 100 μm.

b. RT-PCR analysis of the expression of three germ-line markers in EBs derived from ruminant haSCs.

c. Further immunostaining for three germ-line markers in EBs derived from ruminant haSCs, including ectoderm (NESTIN), mesoderm (BRACHYURY), and endoderm (GATA4). Scale bar, 100 μm.

d. FACS analysis of haploid cells in EBs derived from ruminant haSCs at the indicated time points.

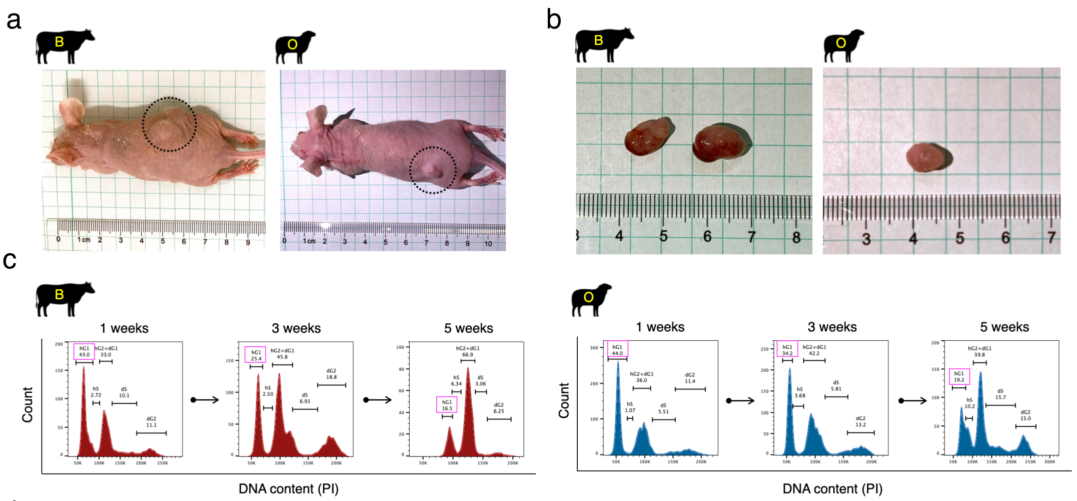


d

| Species | Passage number | haSCs injected into one mouse | Mice age at transplantation | Total mice | Duration | % of teratoma formation | % of 3-germ layer differentiation | Maximum diameter of teratoma (mean ± s.d.; cm) | Teratoma weight (mean ± s.d.; g) | Haploid cells in teratoma (mean ± s.d.; %) |
|---|---|---|---|---|---|---|---|---|---|---|
| B | 25 | $1\times10^9$ | 8 weeks | 3 | 5 weeks | 100 (3/3) | 100 (3/3) | 1.53 ± 0.15 | 1.93 ± 0.25 | 16.67 ± 0.86 |
| | 35 | $1\times10^9$ | 8 weeks | 3 | 5 weeks | 100 (3/3) | 100 (3/3) | 1.50 ± 0.26 | 1.80 ± 0.20 | 16.13 ± 1.46 |
| O | 25 | $1\times10^9$ | 8 weeks | 3 | 5 weeks | 100 (3/3) | 100 (3/3) | 0.80 ± 0.10 | 0.97 ± 0.12 | 19.27 ± 0.60 |
| | 35 | $1\times10^9$ | 8 weeks | 3 | 5 weeks | 100 (3/3) | 100 (3/3) | 0.87 ± 0.21 | 1.07 ± 0.32 | 19.73 ± 1.80 |

### Extended Data Fig. 11

**Ruminant haSCs exhibit competency for the formation of teratoma *in vivo*.**

a. Teratoma produced by injection of ruminant haSCs into the immunodeficient mice.

b. Morphology of teratoma produced by ruminant haSCs.

c. FACS analysis of haploid cells in teratoma derived from ruminant haSCs at the indicated time points.

d. Summary of teratoma formation assay by using ruminant haSCs.

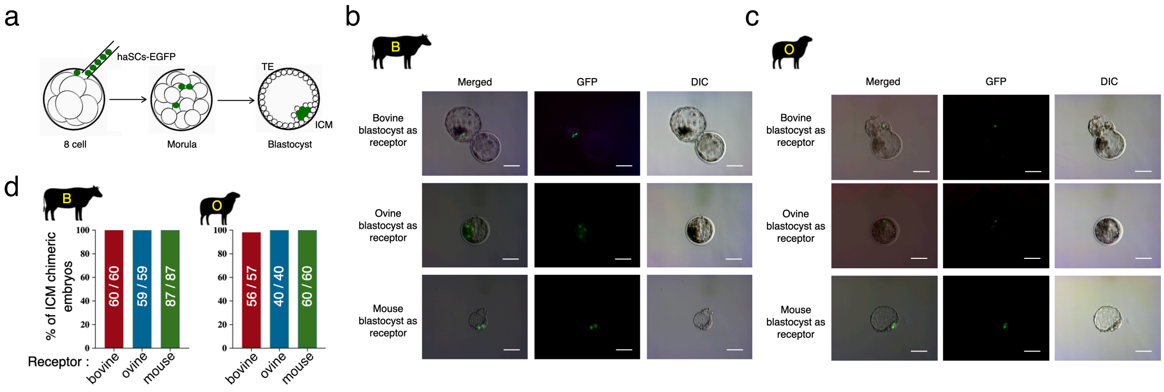


**Extended Data Fig. 12**

**Ruminant haSCs harbour potency for intra- and inter-species chimaeras.**

a. Schematic illustration for producing chimeric blastocysts. Ten EGFP-labeled ruminant haSCs were microinjected into an 8-cell stage embryos, and the chimeric embryos were *in vitro* cultured until the blastocyst stage. ICM, inner cell mass; TE, trophectoderm.

b. Representative phase contrast and fluorescence images showing the incorporation of EGFP-labeled b-haSCs in mouse, bovine, and ovine blastocysts. Scale bar, 50 µm.

c. Representative phase contrast and fluorescence images showing the incorporation of EGFP-labeled o-haSCs in mouse, bovine, and ovine blastocysts. Scale bar, 50 µm.

d. The percentage of chimeric blastocysts with o- or b-haSCs contribution to inner cell mass (ICM). n = 3 independent experiments.

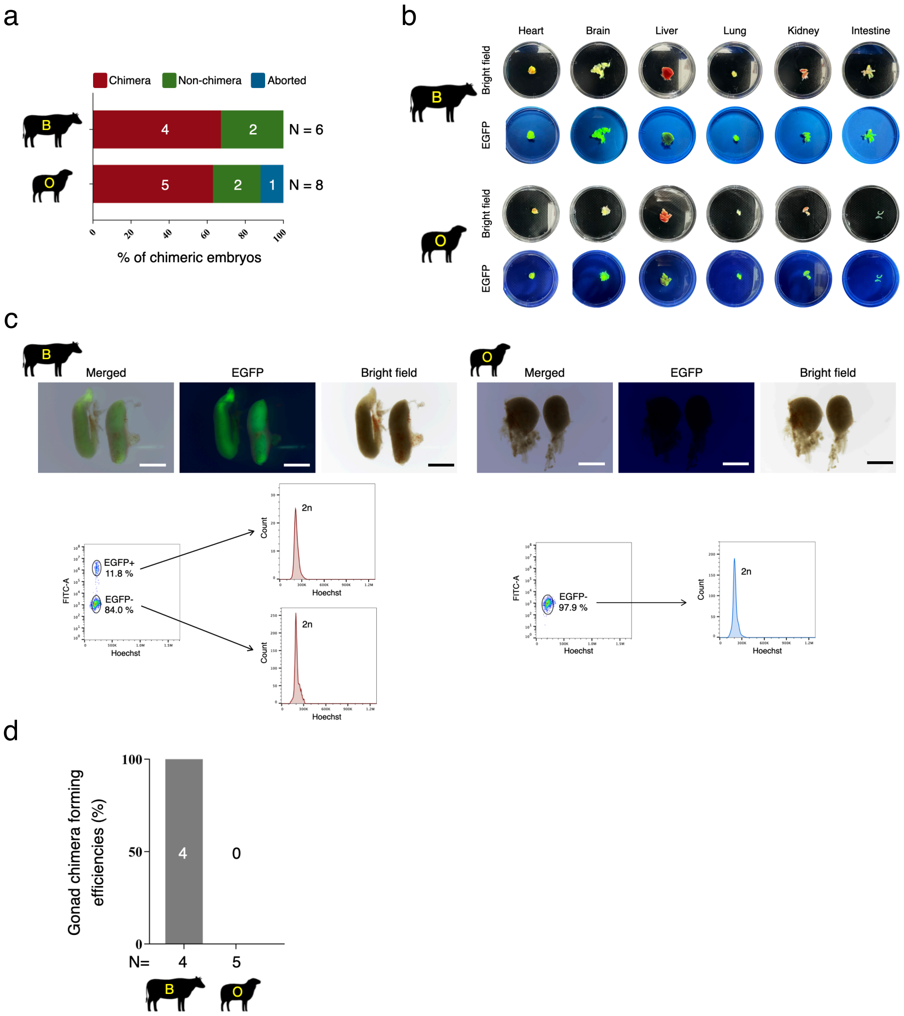


**Extended Data Fig. 13**

**Characterization of intra-species chimaeras derived by b-haSCs or o-haSCs.**

a. Summary of the intra-species chimaeras forming efficiencies. The EGFP-labeled b-haSCs and o-haSCs were microinjected into bovine and ovine 8-cell stage embryos, respectively. The cattle-cattle and sheep-sheep chimeric embryos were transferred into surrogate mothers, and analyzed chimaeras at E40 and E30, respectively. N, the total number of fetuses analyzed for each condition.

b. Representative phase contrast and fluorescence images showing EGFP-labeled b-haSCs or o-haSCs contributed to indicated organs of cattle-cattle and sheep-sheep chimaeras at E40 and E30, respectively.

c. Upper panel: representative phase contrast and fluorescence images showing EGFP-labeled b-haSCs and o-haSCs contributed to the gonads of cattle-cattle and sheep-sheep chimaeras at E40 and E30, respectively. Lower panel: FACS analysis of DNA content of the EGFP-positive and EGFP-negative cells within the chimeric gonads. Note that the EGFP-labeled b-haSCs (but not o-haSCs) contributed to the gonads. Scale bar, 500 μm.

d. Summary of EGFP-labeled b-haSCs or o-haSCs contributed to the gonads in the E40 cattle-cattle and E30 sheep-sheep chimaeras, respectively. N, the total number of fetuses analyzed for each condition.

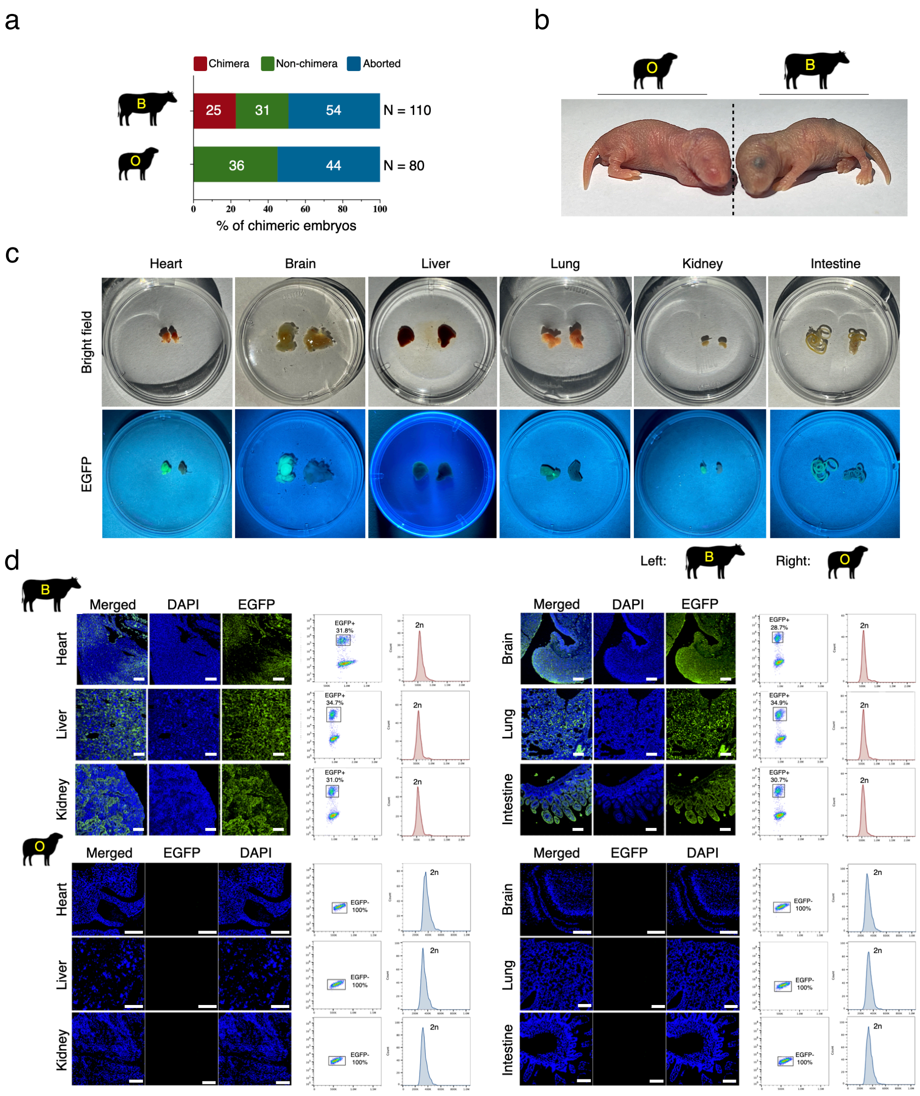


### Extended Data Fig. 14

**Further characterization of cattle-mouse and sheep-mouse chimaeras.**

a. Summary of the inter-species chimaeras forming efficiencies. The EGFP-labeled b-haSCs or o-haSCs were microinjected into 8-cell stage mouse embryos, and the cattle-mouse or sheep-mouse chimeric embryos were transferred into surrogate mouse mothers, and analysed chimaeras at E18.5. N, the total number of fetuses analysed for each condition.

b. Representative images of eye colour in the cattle-mouse and sheep-mouse chimaeras at 2-day-old. The albino mice were used as hosts.

c. Representative phase contrast and fluorescence images showing EGFP-labeled b-haSCs or o-haSCs contributed to indicated organs of cattle-mouse and sheep-mouse chimaeras, respectively.

d. Left panel: immunohistochemistry images showing EGFP-labeled b-haSCs or o-haSCs contributed to different tissues in the cattle-mouse and sheep-mouse chimaeras at 2-day-old. Organ sections were stained with anti-GFP antibody. Right panel: FACS analysis of DNA content of the EGFP-positive or EGFP-negative cells within the different tissues. Scale bar, 200 μm.

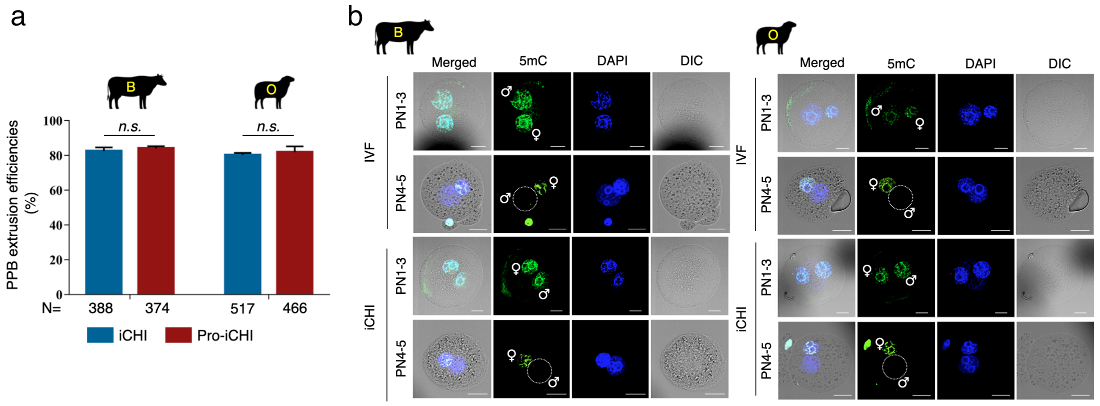


**Extended Data Fig. 15**

**Ruminant haSCs harbour competency for oocyte fertilizing.**

a. Summary of the pseudo-polar body (PPB) extrusion efficiencies in iCHI and Pro-iCHI approaches. Note that the female pronucleus and male pseudo-pronucleus could be efficiently formed in the reconstructed iCHI embryos (110 of 388 in cattle, 110 of 517 in sheep) and Pro-iCHI embryos (110 of 374 in cattle, 110 of 466 in sheep) after extrusion of the PPB, respectively. N, the total number of embryos analyzed for each condition; mean ± s.d.; n > = 3 independent experiments; n.s., not significant by Student's *t*-test.

b. Immunostaining for 5-methylcytosine (5mC) of the bovine and ovine iCHI embryos at the indicated stage. The IVF embryos were used as control. ♀, female pronucleus; ♂, male pronucleus or pseudo-pronucleus; Scale bar, 25 µm.

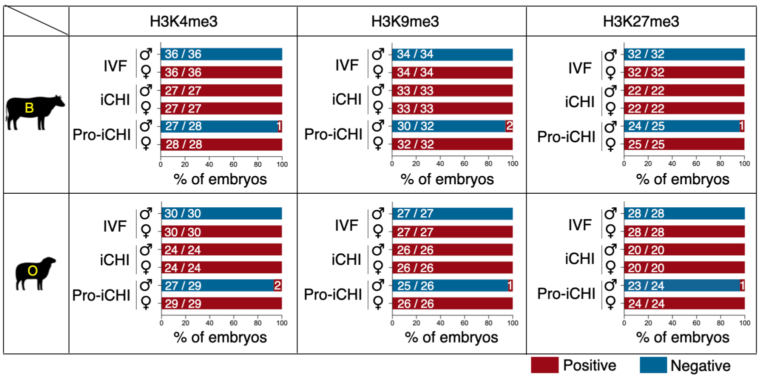


**Extended Data Fig. 16**

**Characterization of histone modification in ruminant embryos.**

Quantifications of H3K4me3, H3K9me3, and H3K27me3 immunostaining analysis in different pronucleus of ruminant iCHI, Pro-iCHI, and IVF embryos. ♀, female pronucleus; ♂, male pronucleus or pseudo-pronucleus.

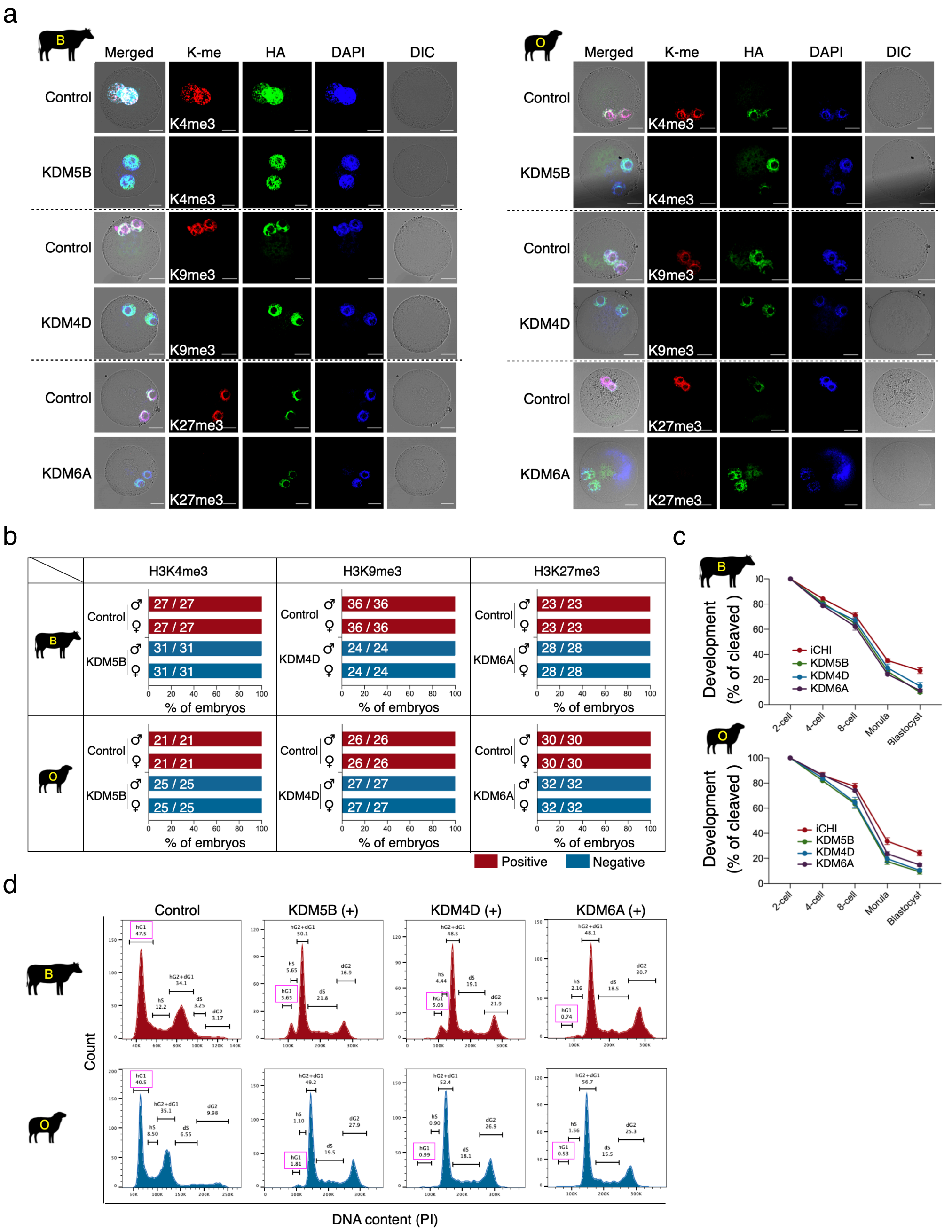


### Extended Data Fig. 17

**Histone demethylases do not amend iCHI-associated symmetric errors.**

a, b. Immunostaining for H3K4me3, H3K9me3, and H3K27me3 of the ruminant iCHI embryos after injection of different mRNA (a). Quantifications of immunostaining results in different embryos (b). The KDM5B, KDM4D, and KDM6A encoding the H3K4me3, H3K9me3, and H3K27me3 demethylases, respectively. The *in vitro* transcription vectors KDMs tagged C-terminally with the hemagglutinin (HA) epitope, which allowed us to track the KDMs proteins in embryos, without the use of specific antibodies. The embryo injected with a blank plasmid vector was used as the negative control. ♀, female pronucleus; ♂, male pronucleus or pseudo-pronucleus; Scale bar, 25 μm.

c. Preimplantation development of ruminant iCHI embryos. The iCHI embryo was injected with mRNA as indicated. The KDM5B, KDM4D, and KDM6A encoding the H3K4me3, H3K9me3, and H3K27me3 demethylases, respectively. mean ± s.d.; n = 3 independent experiments.

d. FACS analysis of haploid cells in ruminant haSCs at 24h after KDM5B, KDM4D, and KDM6A transfection, respectively. The haSCs were transfected with blank plasmid vector was used as the negative control.

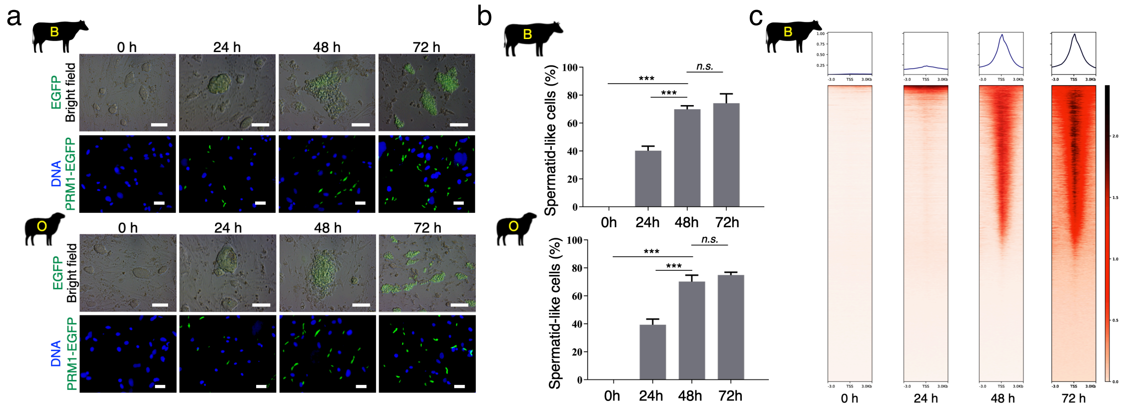


**Extended Data Fig. 18**

**Evaluation of nuclear remodelling efficiency in protaminized ruminant haSCs.**

a. Representative phase contrast and fluorescence images of ruminant haSCs at 0-, 24-, 48-, and 72-h post-protamine-induction (top panel). The bottom panel shows higher-magnification images. Note that after 48h protamine-induction, the cell nuclei acquire a spermatid-like structure, detach from the culture dish, and float in the medium. After 72h of protamine-induction, the protaminized cells showed signs of degeneration. These haSCs harbour the doxycycline (dox)-inducible PRM1-EGFP vector. Scale bars: in top, 100 μm; in bottom, 20 μm.

b. Quantifications of spermatid-like cells in ruminant haSCs at 0-, 24-, 48-, and 72-h post-protamine-induction. mean ± s.d.; n = 3 independent experiments; n.s., not significant; ***$P < 0.001$ by Student's *t*-test.

c. Heatmap and pileup of CUT&Tag signal for the protamine deposition in b-haSCs at 0-, 24-, 48-, and 72-h post-protamine-induction. Each row of the heatmap is a genomic locus and rows are sorted from highest peak intensity to lowest.

a

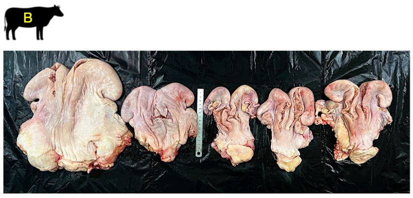


b

| | Items | IVF ( N = 5 ) | Pro-iCHI Growth-normal fetus ( N = 2 ) | Pro-iCHI Growth-retarded fetus ( N = 12 ) |
|---|---|---|---|---|
| Fetus | Fetal body weight (g) | 145.34 ± 4.62 | 144.85 ± 6.58 | 128.27 ± 8.61*** |
| | Fetal eviscerated weight (g) | 101.44 ± 4.05 | 102.90 ± 3.25 | 91.89 ± 5.59** |
| | Crown-rump length (cm) | 11.04 ± 1.13 | 10.05 ± 2.62 | 9.04 ± 1.23** |
| | Umbilical cord diameter (cm) | 1.08 ± 0.22 | 1.00 ± 0.42 | 0.82 ± 0.21* |
| Conceptus | Conceptus length (cm) | 104.00 ± 3.76 | 105.25 ± 5.16 | 106.11 ± 6.50 |
| | Conceptus weight (g) | 1221.76 ± 55.49 | 1224.10 ± 76.93 | 1101.47 ± 46.56*** |
| | Number of cotyledons | 75.92 ± 9.29 | 76.50 ± 4.53 | 75.08 ± 10.20 |
| | Mean cotyledon diameter (cm) | 3.22 ± 0.58 | 3.35 ± 1.63 | 2.31 ± 0.74* |
| | Allantoic fluid (ml) | 338.74 ± 9.34 | 342.85 ± 19.73 | 331.72 ± 4.42* |
| | Amniotic fluid (ml) | 501.18 ± 14.03 | 498.85 ± 17.61 | 472.63 ± 11.89*** |
| | Total placenta weight (g) | 207.16 ± 8.19 | 205.15 ± 11.67 | 192.87 ± 5.75*** |

**Extended Data Fig. 19**

**Characterization of bovine iCHI conceptus.**

a. Representative images of dissected bovine uteri isolated from surrogate mothers after day-65 of iCHI embryo transfer.

b. Summary of the morphometric measurements of bovine conceptuses at E75 produced by Pro-iCHI or IVF techniques. *$P < 0.05$, **$P < 0.01$, ***$P < 0.001$ as compared with IVF group, by two-tailed Student's *t*-test.

a

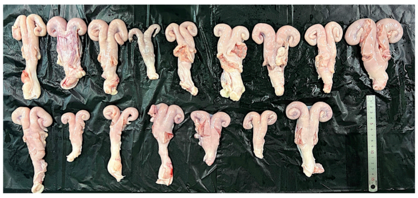

c

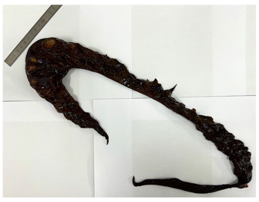

b

| | Items | IVF (N = 5) | Pro-iCHI |
|---|---|---|---|
| | | | Growth-retarded fetus (N = 17) |
| Fetus | Fetal body weight (g) | 153.24 ± 16.13 | 131.45 ± 8.98*** |
| | Fetal eviscerated weight (g) | 101.44 ± 3.81 | 91.98 ± 6.47** |
| | Crown-rump length (cm) | 15.86 ± 4.34 | 11.42 ± 1.92** |
| | Umbilical cord diameter (cm) | 1.52 ± 0.35 | 0.79 ± 0.37*** |
| Conceptus | Conceptus length (cm) | 68.12 ± 5.99 | 67.92 ± 11.38 |
| | Conceptus weight (g) | 1102.06 ± 55.87 | 989.73 ± 29.28*** |
| | Number of cotyledons | 67.08 ± 7.42 | 66.89 ± 9.95 |
| | Mean cotyledon diameter (cm) | 4.18 ± 0.76 | 3.41 ± 0.67* |
| | Allantoic fluid (ml) | 348.18 ± 18.57 | 333.16 ± 12.42* |
| | Amniotic fluid (ml) | 451.18 ± 27.90 | 372.35 ± 32.67*** |
| | Total placenta weight (g) | 298.68 ± 16.46 | 201.16 ± 27.17*** |

**Extended Data Fig. 20**

**Characterization of ovine iCHI conceptus.**

a. Representative images of dissected ovine uteri isolated from surrogate mothers after day-65 of iCHI embryo transfer.

b. Summary of the morphometric measurements of ovine conceptuses at E65 produced by Pro-iCHI or IVF techniques. $^{*}P < 0.05$, $^{**}P < 0.01$, $^{***}P < 0.001$ as compared with IVF group, by two-tailed Student's *t*-test.

c. An image of stillborn ovine Pro-iCHI conceptus isolated from surrogate mother after day-150 of embryo transfer.

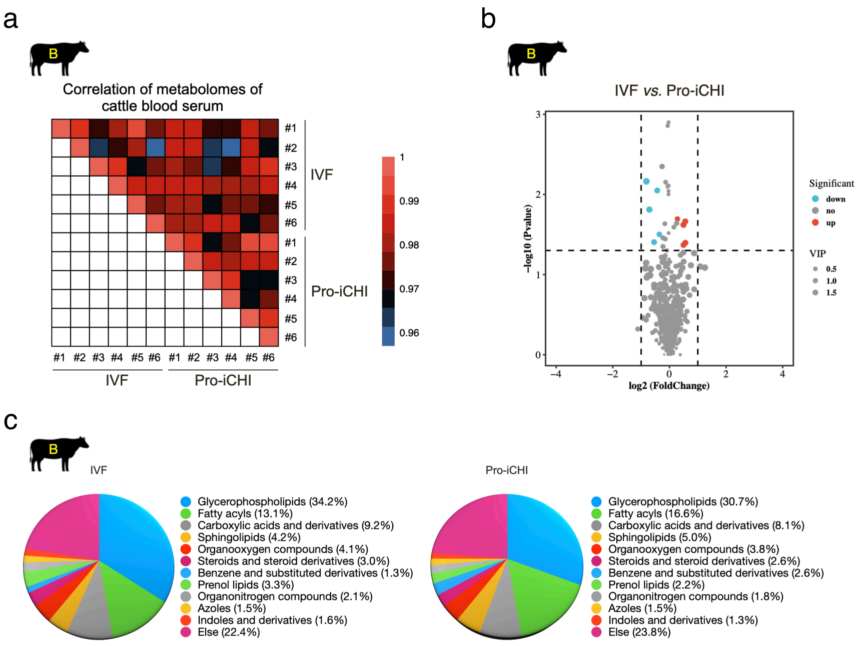


**Extended Data Fig. 21**

**Pro-iCHI and IVF live cattle have similar serum metabolomes.**

a. Heatmap showing the correlation of blood serum metabolomes of Pro-iCHI and IVF cattle in different replicates at 18-month-old.

b. Volcano plot showing the metabolic changes between Pro-iCHI and IVF cattle at 18-month-old. The vertical and horizontal dotted lines show the cut-off of fold-change = ± 2, and *p*-value = 0.05, respectively.

c. Pie charts showing the metabolite class composition in Pro-iCHI and IVF cattle at 18-month-old.

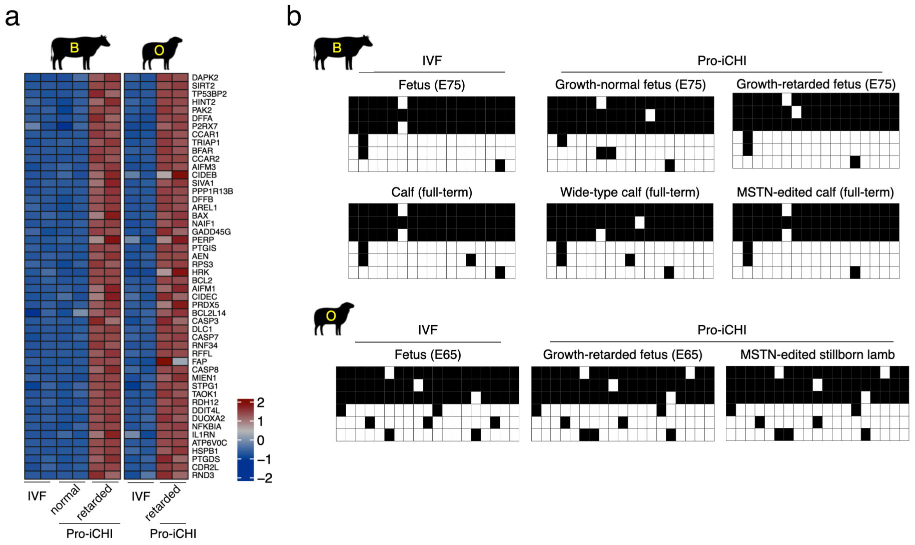


**Extended Data Fig. 22**

**Transcriptome and methylation analysis of ruminant haSCs produced animals.**

a. Heatmap showing cross-species comparison of orthologous genes in bovine and ovine fetuses produced by Pro-iCHI or IVF approach. Note that the expression level of apoptosis and degeneration-related genes was significantly increased (FC > 2, FPKM > 2) in growth-retarded Pro-iCHI fetuses compared with normal Pro-iCHI fetuses and control IVF fetuses. FC, fold change; FPKM, fragments per kilobase of transcript per million mapped reads.

b. Bisulfite pyrosequencing analysis of paternally imprinted H19 DMR in bovine and ovine fetuses produced by Pro-iCHI or IVF approach. Note that all the fetuses maintained allelic-biased DNA methylation in H19 DMR. The filled and open squares represent methylated and unmethylated CpG sites, respectively.

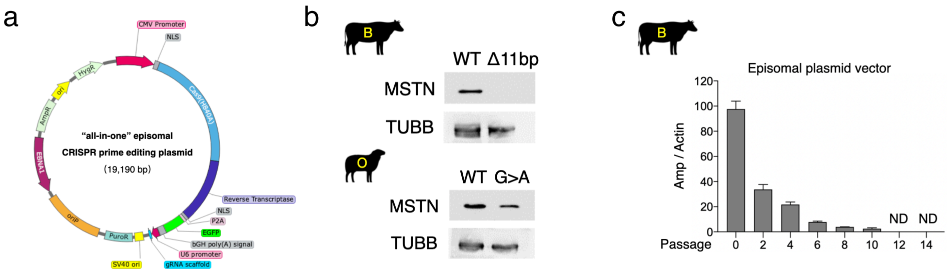


**Extended Data Fig. 23**

**Establishment of an integration-free ePE system to generate gene-edited haSCs.**

a. Schematic representation of the ePE vector used in this study. This “all-in-one” episomal plasmid contained the necessary elements of CRISPR-prime editor (PE), including Cas9-nickase and reverse-transcriptase (RT) fusion sequence, and a prime-editing guide RNA (pegRNA). The vector also contains an EF1a promoter-driven EGFP for tracking transfection efficiency. Puro, Puromycin resistance gene.

b. Western blots analysis of MSTN protein in wild-type (WT) and MSTN-edited ruminant haSCs.

c. Episomal plasmid was decreased within haSCs over time after withdrawing puromycin drug selection. Numbers in the X-axis indicate cell passage number (passage cells every 5 days). Amp, ampicillin; Actin, beta-actin; mean ± s.d., n = 3 independent experiments; amplification cycle was maintained at 40 and the cycle threshold (Ct) values > 35 were considered as not detected (ND).

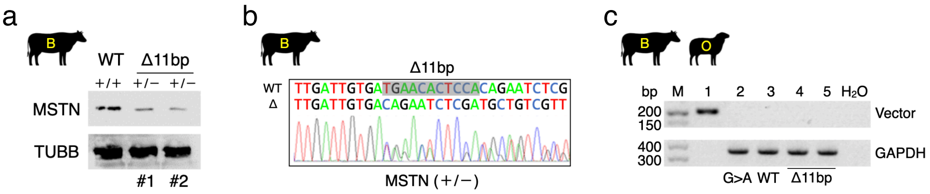


**Extended Data Fig. 24**

**Rapid generation of gene-edited livestock using the Pro-iCHI approach.**

a. Western blots analysis of MSTN protein in wild-type (WT) and MSTN-edited live cattle.

b. Sanger sequencing results of the targeting site in wild-type (WT) and MSTN-edited live cattle, the deletion sizes (Δ) are indicated.

c. Genomic PCR showing the absence of exo-vector in lamb and calf produced by Pro-iCHI::ePE system. The positive and negative controls were amplified from the episomal plasmid and water, respectively. Lane 1: positive control; Lane 2: sheep; Lanes 3-5: cattle; M, DNA marker.
