## Supplementary information for "One-step generation of modified cattle and sheep from spermatid-like haploid stem cells"

6 Deceased

*

**This file includes:**

Supplementary Note (Page 2-3)

Supplementary Figures 1-8 (Page 4-11)

Supplementary Table 1-8 (Page 12-27)

#### Supplementary Note

To determine whether b-haSCs harbour the competency for primordial germ cells (PGC) induction *in vitro*, we first generated the dual reporter b-haSCs lines, which contained two PGC-specific reporters: PRDM1 (also known as BLIMP1)-tdTomato (PT) and TFAP2C-mNeonGreen (TG; Supplementary Fig. 1a-d). Following a recently established bovine protocol (Shirasawa A., *et al.*, J Reprod Dev. 2024, PMID: 38355134), we successfully differentiated b-haSCs into PGC-like cells (Supplementary Fig. 2a). Particularly, the number of $PT^{+}$::$TG^{+}$ positive cells continued to increase during the PGC-induction (Fig. 3e). FACS analysis revealed that 31%, 65%, and 73% of cells were $PT^{+}$::$TG^{+}$ after 2, 3, and 4 Days of induction, respectively (Fig. 3f). We also found that no haploid cells could be detected during the PGC-induction (Supplementary Fig. 2b).

We also performed the bulk RNA-seq (buRNA-seq) analysis and found that the expression levels of critical PGC-markers, such as PRDM1/BLIMP1, SOX17, TFAP2C, NANOS3, and KIT, were upregulated during PGC-induction (Supplementary Fig. 3a,b). Specifically, SOX2, which is upregulated in rodent PGCs but not in other mammalian PGCs, was downregulated during PGC induction from b-haSCs (Irie N., *et al.*, Cell. 2015, PMID: 25543152; Yamaji M., *et al.*, Nat Genet. 2008, PMID: 18622394; Shirasawa A., *et al.*, J Reprod Dev. 2024, PMID: 38355134; Yu L., *et al.*, Cell Stem Cell. 2021, PMID: 33271070). To dissect the cell population at higher resolution, we further conducted single-cell RNA sequencing (scRNA-seq). Consistent with buRNA-seq analysis, 86.6% of the whole cell population expressed PGC-marker genes at the end stage of PGC-induction (Supplementary Fig. 3c).

In addition to the transcriptome analysis, immunofluorescence (IF) assay confirmed that the $PT^{+}$::$TG^{+}$ cells co-expressed endogenous PRDM1/BLIMP1 and TFAP2C proteins, as well as SOX17, OCT4, and NANOG, but not SOX2 (Supplementary Fig.

4a). Consistent with a recent report (Shirasawa A., *et al.*, J Reprod Dev. 2024, PMID: 38355134), we found that DNA methylation (5-methylcytosine, 5mC) and H3K9me2 levels decreased, while the H3K27me3 level increased in $PT^{+}$::$TG^{+}$ positive cells when compared to the negative cells (Supplementary Fig. 4b). Taken together, our results demonstrate that, in addition to intra-/inter-species chimera competency, b-haSCs are also amenable to direct PGC induction *in vitro*.

#### Supplementary Figures

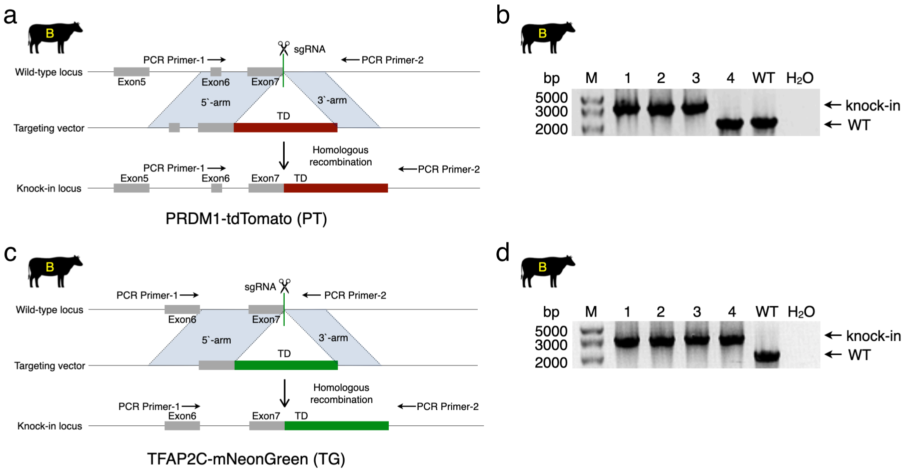

**Supplementary Fig. 1 l Generation of PGC-reporter b-haSC lines.**

(a). Strategy to introduce the tdTomato reporter into the PRDM1/BLIMP1 locus using CRISPR/Cas9 via homology-directed repair (HDR). The green line indicates the location of the sgRNA used to target the PRDM1/BLIMP1 locus. Arrows indicate the binding sites of PCR primers used to confirm the insertion of tdTomato into the PRDM1/BLIMP1 locus.

(b). Confirmation of tdTomato insertion using PCR genotyping. Three of four b-haSC lines tested showed correct insertion of tdTomato into PRDM1/BLIMP1 locus. The primer binding sites are depicted in (a). Wild type (WT) haSCs are used as controls; M, DNA marker.

(c). Strategy to introduce the mNeonGreen reporter into the TFAP2C locus using CRISPR/Cas9 via HDR. The green line indicates the location of the sgRNA used to target the TFAP2C locus. Arrows indicate the binding sites of PCR primers used to confirm the insertion.

(d). Confirmation of mNeonGreen insertion using PCR genotyping. All tested b-haSC lines showed the correct insertion of mNeonGreen into the TFAP2C locus. The primer binding sites are depicted in (c). Wild type (WT) haSCs are used as controls; M, DNA marker.

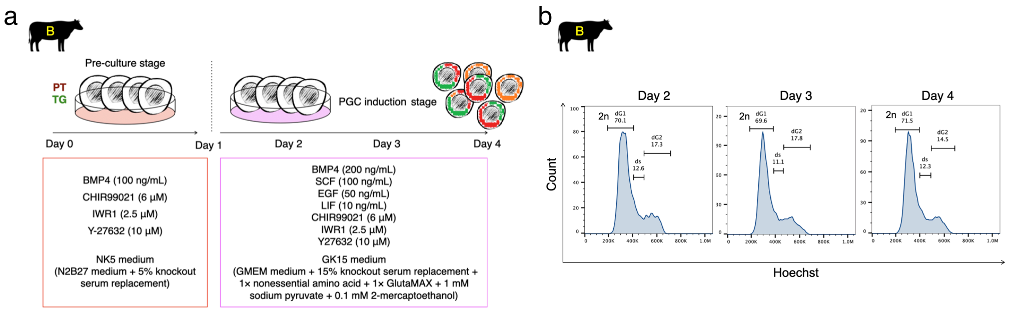

**Supplementary Fig. 2 | *In vitro* induction of PGCs from b-haSCs.**

(a). Schematic overview of direct induction of PGCs from b-haSCs, according to the recently established approach (Shirasawa A., *et al.*, J Reprod Dev. 2024, PMID: 38355134).

(b). Fluorescence-activated cell sorting (FACS) analysis of haploid cells during the induction of PGCs from b-haSCs. dG1, the diploid G1 phase of the cell cycle; dS, the diploid S phase of the cell cycle; dG2, the diploid G2 phase of the cell cycle.

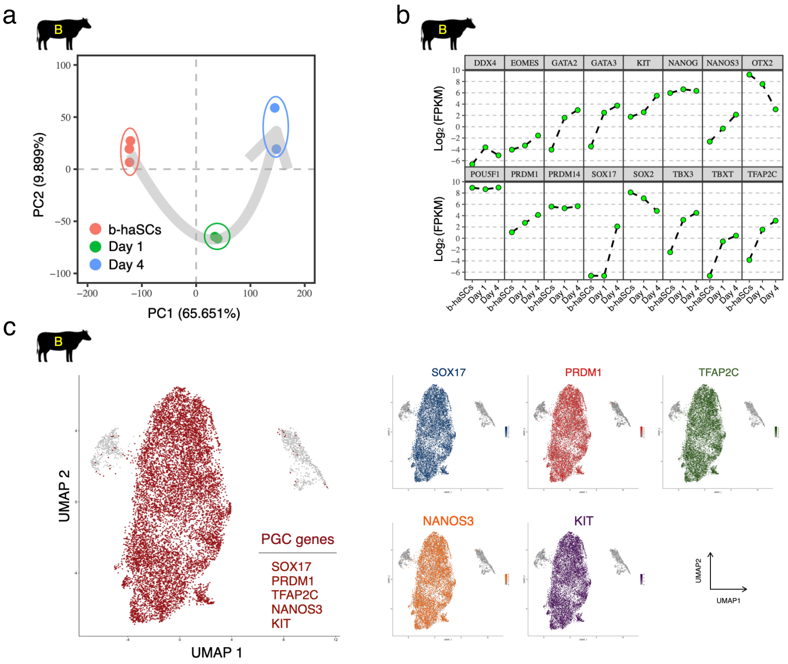

**Supplementary Fig. 3 l Transcriptome analysis of PGC-induction from b-haSCs.**

(a). Principal-component analysis (PCA) of PGCs induction from b-haSCs using buRNA-seq data, samples were coloured by time points.

(b). Expression dynamics of selected PGC-marker genes during induction of PGCs from b-haSCs by buRNA-seq. Each dot represents one biological replicate. The expression levels are represented by fragments per kilobase per million (FPKM).

(c). Expression patterns of selected PGC-marker genes in the PGCs induced from b-haSCs at the single-cell level by scRNA-seq.

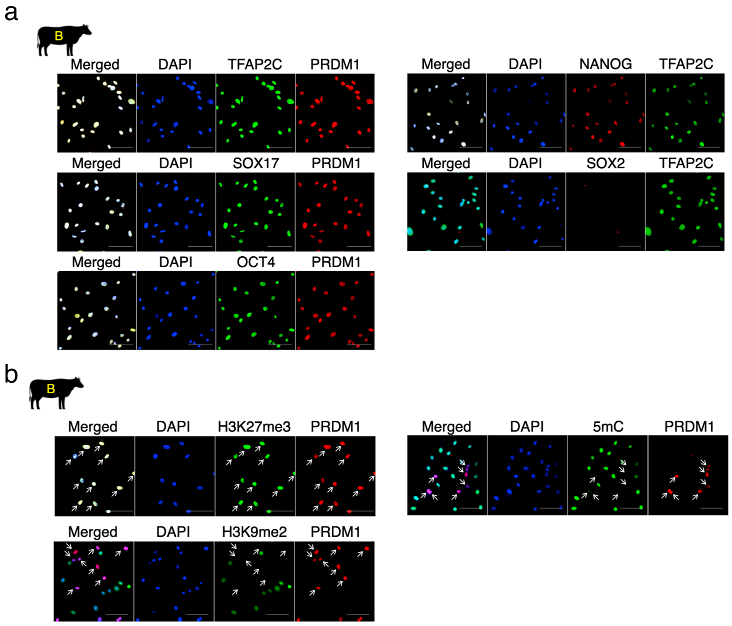

**Supplementary Fig. 4 l Further characterization of PGCs induced from b-haSCs.**

(a). Representative IF images showing the expression of PGC-markers in the PGCs induced from b-haSCs, including PRDM1/BLIMP1, TFAP2C, SOX17, OCT4, and NANOG (but not SOX2). Scale bar, 100 µm.

(b). Representative IF images of epigenomic markers in the PGCs induced from b-haSCs, including 5mC, H3K9me2, and H3K27me3. Arrows, delineate PRDM1-positive cells. Scale bar, 100 µm.

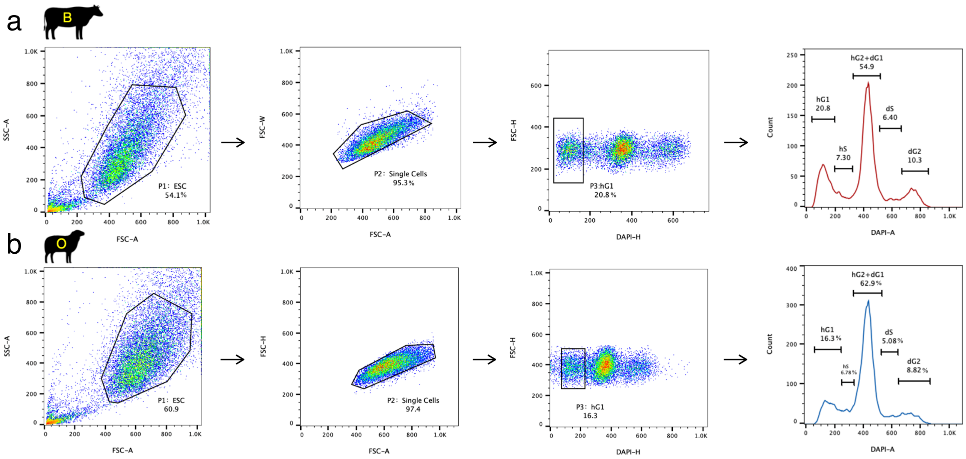

**Supplementary Fig. 5 I FACS enrichment of haploid cells.**

FACS gating examples for bovine (a) and ovine (b) haSCs, according to the published protocols (details in Methods). The live cells are stained with Hoechst 33342 and gate P1 is centered around the bulk of the SCs. The cell clumps are removed by setting gate P2 around the main cell population. The cells gated on P2 are subsequently sorted using gate. hG1, the haploid G1 phase of the cell cycle; hS1, the haploid S phase of the cell cycle; hG2, the haploid G2 phase of the cell cycle; dG1, the diploid G1 phase of the cell cycle; dS, the diploid S phase of the cell cycle; dG2, the diploid G2 phase of the cell cycle. These images are representative examples of the FACS sorting shown in Fig. 1c.

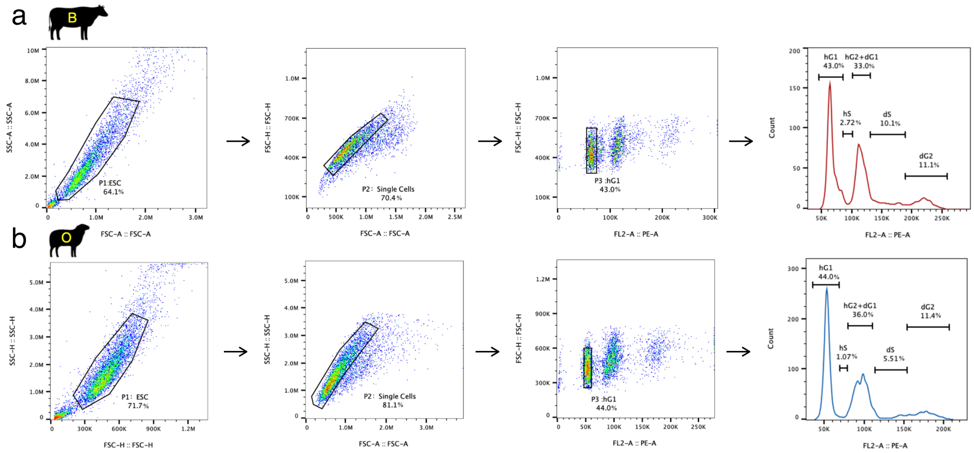

**Supplementary Fig. 6 l FACS analysis of haSCs.**

The FACS analysis of bovine (a) and ovine (b) haSCs follows a setup similar to that of the haploid cell sorting (details in Supplementary Fig. 1 legend). The samples were fixed and stained with propidium iodide (PI) as described in the Methods. hG1, the haploid G1 phase of the cell cycle; hS1, the haploid S phase of the cell cycle; hG2, the haploid G2 phase of the cell cycle; dG1, the diploid G1 phase of the cell cycle; dS, the diploid S phase of the cell cycle; dG2, the diploid G2 phase of the cell cycle. These images are representative examples of the FACS analysis shown in Extended data Fig. 10d.

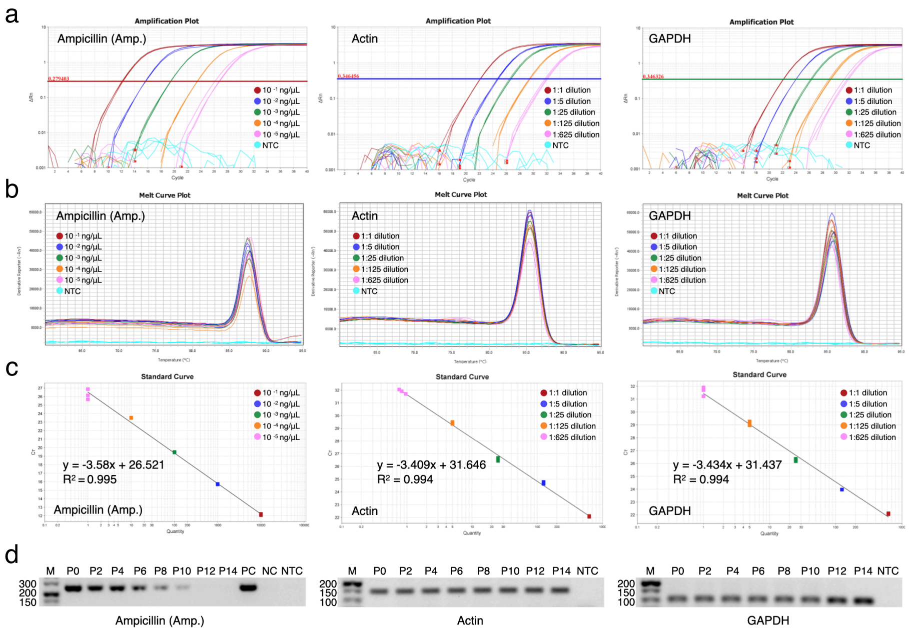

**Supplementary Fig. 7 | Relevant information for real-time PCR experiments.** Representative plots of amplification (a), melt curves (b), standard curves (c) and gel visualization (d) of RT-qPCR are shown in Extended Data Fig. 23e. Each line or dot represents one experiment. In (c), the solid line is the fitted line and $R^2$ is the Pearson's correlation coefficient. In (d), positive control (PC) was amplified from the plasmid; Negative-control (NC) was amplified from the non-transfected wild-type (WT) cells; No template control (NTC) was amplified from the water ($H_2O$); M, DNA marker; P, cell passage.

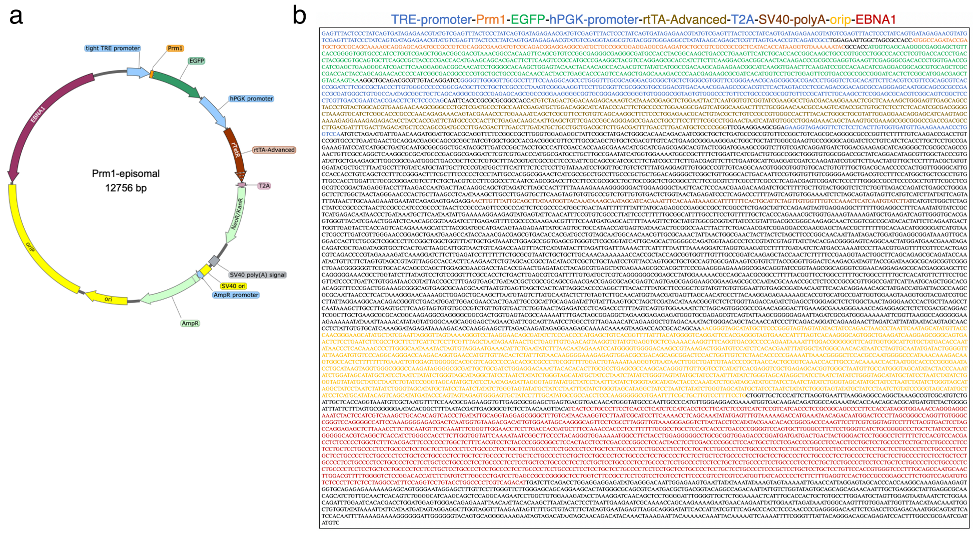

**Supplementary Fig. 8 I Sequence information of PRM1-EGFP plasmid.**

Circular map (a) and sequence information (b) of episomal-based doxycycline (dox)-inducible PRM1-EGFP plasmid. Prm1, protamine1 gene; EGFP, enhanced green fluorescent protein gene; tight TRE promoter, tight tetracycline-responsive element promoter; hPGK promoter, human phosphoglycerate kinase promoter; rtTA-Advanced, an improved version of the reverse tetracycline-controlled transactivator protein; T2A, 2A regions of thosea asigna virus; NeoR/KanR, aminoglycoside phosphotransferase from Tn5 confers resistance to kanamycin in bacteria and neomycin and G418 in eukarytoes; SV40 ori, simian vacuolating virus 40 origin of replication; SV40 poly(A) signal, simian vacuolating virus 40 poly(A) signal; AmpR, ampicillin resistance gene; ori, origin of replication; orip, epstein-bar virus latent origin of replication; EBNA1, EBV nuclear antigen-1 replication transactivator of EBV.

#### Supplementary Tables

**Table 1. Developmental rates of ruminant embryos derived using different methods.**

| Species | Group | Method | No. of replicates | No. of cultured | No. of 2-cell (% of cultured) | No. of 4-cell (% of cleaved) | No. of 8-cell (% of cleaved) | No. of morula (% of cleaved) | No. of blastocyst (% of cleaved) |
|---|---|---|---|---|---|---|---|---|---|
| 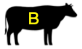  | Diploid embryos | IVF | 3 | 154 | 138 (89.61) | 123 (89.13) | 115 (83.33) | 99 (71.74) | 95 (68.84) |
| | Haploid embryos | Injecting sperm | 5 | 367 | 276 (75.20)*** | 234 (84.78)* | 209 (75.72)** | 119 (43.12)*** | 86 (31.16)*** |
| | | Removing pronuclei | 5 | 314 | 235 (74.84)*** | 193 (82.13)* | 172 (73.19)** | 94 (40.00)*** | 71 (30.21)*** |
| 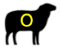  | Diploid embryos | IVF | 3 | 517 | 404 (78.14) | 353 (87.38) | 332 (82.18) | 252 (62.38) | 217 (53.71) |
| | Haploid embryos | Injecting sperm | 3 | 614 | 366 (59.61)*** | 286 (78.14)** | 244 (66.67)*** | 133 (36.34)*** | 94 (25.68)*** |
| | | Removing pronuclei | 3 | 584 | 351 (60.10)*** | 265 (75.50)** | 230 (65.53)*** | 119 (33.90)*** | 89 (25.36)*** |

*$P < 0.05$, **$P < 0.01$, ***$P < 0.001$ as compared with IVF group, by two-tailed Student's *t*-test.

**Table 2. Developmental rates of ruminant embryos derived using different iCHI approaches.**

| Species | Group | Treated with | | No. of replicates | No. of reconstructed embryos | % 1-cell with PPB per reconstructed embryos ± S.D. | % 2-cell per 1-cell with PPB ± S.D. | % 4-cell per 2-cell ± S.D. | % 8-cell per 2-cell ± S.D. | % morula per 2-cell ± S.D. | % blastocyst per 2-cell ± S.D. |
|---|---|---|---|---|---|---|---|---|---|---|---|
| 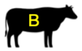 | iCHI | --- | | 3 | 388 | 82.58 ± 2.04 | 87.31±0.90 | 84.21±1.61 | 70.98±2.40 | 35.10±1.37 | 27.12±2.42 |
| | mRNA-iCHI | KDM5B # | | 3 | 219 | 83.84 ± 0.68 | 82.15±0.34*** | 80.94±0.77* | 64.53±1.90* | 26.58±0.62*** | 9.89±0.90*** |
| | | KDM4D ## | | 3 | 261 | 83.94 ± 1.00 | 79.91±0.65*** | 79.50±1.12* | 66.97±2.73 | 29.33±2.92* | 14.80±2.81** |
| | | KDM6A ### | | 3 | 253 | 84.66 ± 1.09 | 81.38±0.70*** | 78.74±0.51** | 62.14±2.87* | 24.19±0.72*** | 11.33±1.19*** |
| | Pro-iCHI | Protamine | 24h | 5 | 362 | 83.69 ± 2.27 | 88.43 ± 0.50 | 85.11 ± 1.44 | 76.94 ± 2.18* | 58.74 ± 1.87*** | 51.97 ± 1.30*** |
| | | | 48h | 5 | 374 | 84.03 ± 1.22 | 89.26±1.12* | 86.73±2.23 | 80.82±1.64*** | 69.72±2.37*** | 64.91±3.34*** |
| 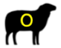 | iCHI | --- | | 3 | 517 | 80.22 ± 1.26 | 61.18±0.31 | 86.15±2.35 | 77.37±2.40 | 33.68±2.75 | 24.22±2.21 |
| | mRNA-iCHI | KDM5B # | | 3 | 295 | 80.48 ± 1.45 | 60.18±2.34 | 81.98±1.90 | 63.43±3.64** | 17.43±2.23** | 9.32±1.93*** |
| | | KDM4D ## | | 3 | 338 | 81.01 ± 2.49 | 59.46±0.74* | 84.10±1.91 | 64.66±4.05** | 19.69±2.96** | 10.46±1.15*** |
| | | KDM6A ### | | 3 | 307 | 78.95 ± 1.92 | 58.92±1.24* | 86.55±1.29 | 74.10±0.92 | 23.70±1.57** | 14.78±1.07** |
| | Pro-iCHI | protamine | 24h | 5 | 400 | 79.86 ± 1.14 | 70.01 ± 1.28*** | 86.46 ± 0.85 | 78.65 ± 2.47 | 51.72 ± 3.48*** | 41.80 ± 1.13*** |
| | | | 48h | 5 | 466 | 81.89 ± 3.33 | 75.37±2.05*** | 87.68±1.88 | 80.20±3.86 | 62.88±2.29*** | 51.22±1.78*** |

*$P < 0.05$, **$P < 0.01$, ***$P < 0.001$ as compared with iCHI group, by two-tailed Student's *t*-test; PPB, pseudo-polar body.

### KDM5B (lysine[K]-specific demethylase 5B), specifically reduces the level of H3K4me3; ## KDM4D (lysine[K]-specific demethylase 4D), specifically reduces the level of H3K9me3; ### KDM6A (lysine[K]-specific demethylase 6A), specifically reduces the level of H3K27me3.

**Table 3. Development of ruminant embryos derived by iCHI and Pro-iCHI.**

| Species | Group | No. of reconstructed embryos with pseudo-polar body (1-cell) | No. of 2-cell (% of 1-cell) | No. of blastocyst (% of 2-cell) | Embryos transferred stage | No. of embryos transferred (ET) per recipient | No. of recipients | No. of fetuses at embryonic day 65 (E65) | No. of fetuses at embryonic day 75 (E75) | No. of pups at full-term (% of ET) |
|---|---|---|---|---|---|---|---|---|---|---|
| | iCHI | 205 | 179(87.32) | 49(27.37) | blastocyst | 1 | 20 | 0 | --- | 0 |
| 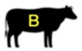 | Pro-iCHI | 71 | 63(88.73) | 41(65.08) | blastocyst | 1 | 14 | --- | 2 normal / 12 retarded | --- |
| | | 51 | 45 (88.24) | 29 (64.44) | blastocyst | 1 | 7 | --- | --- | 1 (14.29) |
| | iCHI | 361 | 221(61.22) | --- | 2-cell | 8 | 26 | 0 | --- | --- |
| 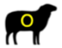 | Pro-iCHI | 249 | 188(75.50) | --- | 2-cell | 8 | 20 | 0 normal / 17 retarded | --- | --- |
| | | 122 | 92 (75.40) | --- | 2-cell | 8 | 9 | --- | --- | 0 (0) |

**Table 4. Development of MSTN-edited ruminant derived by Pro-iCHI.**

| Species | Edited type | MSTN loci | No. of reconstructed embryos with pseudo-polar body (1-cell) | No. of 2-cell (% of 1-cell) | No. of blastocyst (% of 2-cell) | Embryos transferred stage | No. of embryos transferred (ET) per recipient | No. of recipients | No. of pups at full-term (% of ET) |
|---|---|---|---|---|---|---|---|---|---|
| 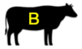 | deletion | Δ 11 bp | 66 | 58 (87.88) | 37 (63.79) | blastocyst | 1 | 15 | 2 (13.33) |
| 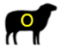 | mutation | G > A | 362 | 272 (75.14) | --- | 2-cell | 8 | 32 | 1 (0.39) |

**Table 5. List of antibodies used in this study.**

| Target | Source | Catalog number | Application |
|---|---|---|---|
| OCT4 | Santa Cruz | sc-5279 | IF (1:200)<br>WB (1:500) |
| SOX2 | Santa Cruz | sc-365823 | IF (1:200) |
| OTX2 | R&D Systems | AF1979 | IF (1:250)<br>WB (1:500) |
| CDX2 | BioGenex | AM392-5M | IF (1:300) |
| NANOG | Thermo | 14-5768-82 | IF (1:200) |
| NESTIN | Santa Cruz | sc-21247 | IF (1:100) |
| TUJ1 | R&D Systems | MAB1195 | IF (1:500) |
| BRACHYURY | Santa Cruz | sc-17745 | IF (1:200) |
| SMA | Abcam | ab5694 | IF (1:200) |
| GATA4 | Santa Cruz | sc-1237 | IF (1:100) |
| FOXA2 | Abnova | H00003170-M10 | IF (1:500) |
| PRM1 | Proteintech | 15697-1-AP | CUT&Tag (1:50) |
| SOX17 | R&D Systems | AF1924 | IF (1:100) |
| H3K4me3 | Abcam | ab8580 | IF (1:50)<br>CUT&Tag (1:50) |
| H3K27me3 | Cell Signaling | 9733S | IF (1:50)<br>CUT&Tag (1:50) |
| H3K9me3 | Abcam | ab176916 | IF (1:50) |
| H3K9me2 | Millipore | 07-441 | IF (1:500) |
| 5mC | Abcam | ab214727 | IF (1:50) |
| 5hmC | Active Motif | 40900 | IF (1:50) |

| | | | |
|---|---|---|---|
| PRDM1 | Thermo | 14-5963-82 | IF (1:100) |
| TFAP2C | R&D systems | AF5059 | IF (1:500) |
| DNMT3A | Proteintech | 20954-1-AP | WB (1:500) |
| DNMT3B | Proteintech | 26971-1-AP | WB (1:500) |
| MSTN | Proteintech | 19142-1-AP | WB (1:300) |
| GAPDH | Proteintech | 10494-1-AP | WB (1:1000) |
| AlexaFluor 594 goat anti-rabbit IgG | Thermo | A-11037 | IF (1:1000) |
| AlexaFluor 488 goat anti-rabbit IgG | Thermo | A-11034 | IF (1:1000) |
| AlexaFluor 594 goat anti-mouse IgG | Thermo | A-11032 | IF (1:1000) |

**Table 6. List of primers and sgRNAs used in this study.**

| Genes | Application | Species | Sequences (5'-3') | Position | GenBank accession number |
|---|---|---|---|---|---|
| DNMT3A | RT-PCR | 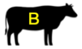  | forward: GGTGGAAAGCAGTGACACACC<br>reverse: CCTTGGCTTTCTTCTCAGCCT | 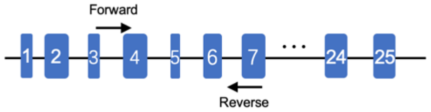  | NM_001206502 |
| | | 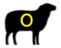  | forward: GGTGGAAAGCAGTGACACACC<br>reverse: CCTTGGCTTTCTTCTCAGCCT | 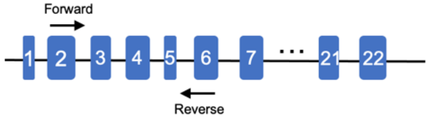  | XM_027966565 |
| | | 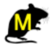  | forward: GTCTCAGCCCATGGCCCAG<br>reverse: CTCTCCAGAGGCCTGGTTCTC | 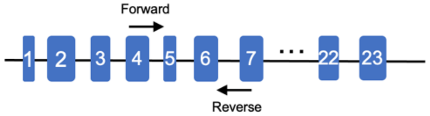  | NM_001421857 |
| DMNT3B | RT-PCR | 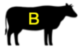  | forward: ACAAGCACGCCAACAGAAGG<br>reverse: CGTGCCTGCAGAGGATCCC | 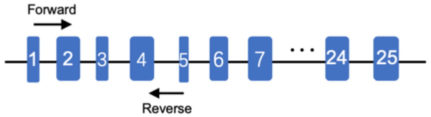  | NM_181813 |
| | | 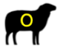  | forward: GAGGCCATCAGCACCCCTG<br>reverse: CCAGGCGGGTATAGGGCGT | 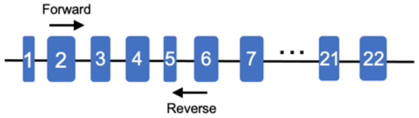  | XM_042230025 |
| | | 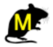  | forward: GAAGAAGAGGGTGCCAGCGG<br>reverse: TGAAGATGATGCTCGACGTCT | 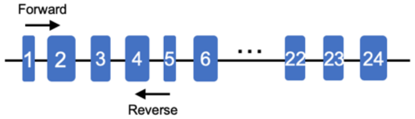  | NM_001003961 |
| OTX2 | RT-PCR | 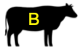  | forward: CGCCTTATGCAGTCAATGGGC<br>reverse: GGCTGGGACTGAGGTGCTA | 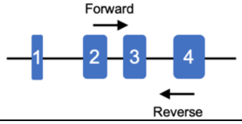  | NM_001193201 |
| | | 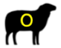  | forward: GGCTGAGTCTGACCACTTCGG<br>reverse: GAGCACTGCTGCTGGAAATGG | 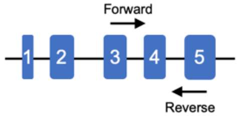  | XM_015097088 |

| | | | | | |
|---|---|---|---|---|---|
| | | M | forward: TCTAAAGCAACCGCCTTACGC | 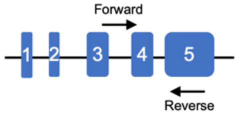  | NM_001286481 |
| | | | reverse: TGGCCACTTGTTCCACTCTCT | | |
| OCT4 | RT-PCR | B | forward: ACCTTGAACAATTTGCCAAGC | 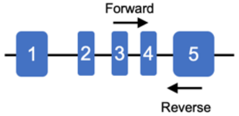  | NM_174580 |
| | | | reverse: GCTAATTTGCTGCAGGGTGG | | |
| | | O | forward: GCAGGCCCGAAAGAGAAAGC | 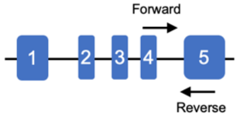  | XM_042237451 |
| | | | reverse: CCTCACCCTCAGGGAATGGG | | |
| | | M | forward: GAGGATCACCTTGGGGTACAC | 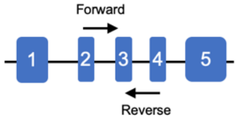  | NM_013633 |
| | | | reverse: CATCCTTCTCTAGCCCAAGCT | | |
| GAPDH | RT-PCR | B | forward: CGGAGTGAACGGATTCGGC | 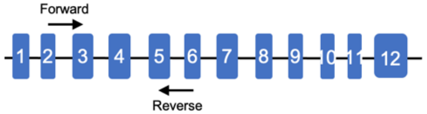  | NM_001034034 |
| | | | reverse: GGTGCAGAGATGATGACCCTC | | |
| | | O | forward: TGGACATCGTTGCCATCAATG | 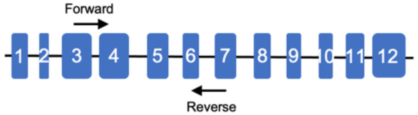  | NM_001190390 |
| | | | reverse: GGAGGCATTGCTGACAATCTT | | |
| | | M | forward: GAGTGTTTCCTCGTCCCGTAG | 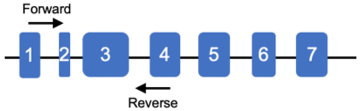  | NM_001289726 |
| | | | reverse: GACCCTTTTGGCTCCACCCT | | |
| NESTIN | RT-PCR | B | forward: CCACTGAGCAGTTCCAGCTG | 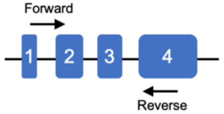  | NM_001206591 |
| | | | reverse: CAGAAAGGTTGGCACAGGTG | | |
| | | O | forward: GCGGACCACTGAGCAGTTC | 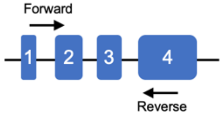  | XM_012183418 |
| | | | reverse: AGGTTGGCACAGGTGTTTCC | | |
| GATA4 | RT-PCR | B | forward: GAAGCTCCATGGCGTCCCC | | NM_001192877 |

| | | | | | |
|---|---|---|---|---|---|
| | | | reverse: GGCTGACAGAAGACGCGTAG | 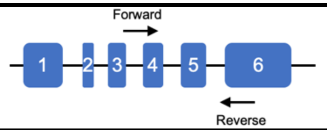 | |
| | | O | forward: GGCTCCTACTCCAGCCCCTAC<br>reverse: CAGTTGGCGCAGGAAAGGC | 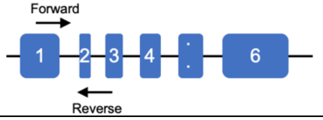 | XM_060411039 |
| AFP | RT-PCR | B | forward: GAAGGCACCCTGTCCTGTATGC<br>reverse: CTCTTTCCCCATCCTGCAGAC | 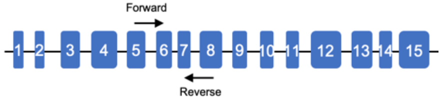 | NM_001034262 |
| | | O | forward: CCCACTGGCAGTGAGCAACCTG<br>reverse: GCATTCGGTGGCATTTTCAGC | 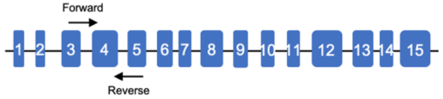 | XM_060417195 |
| FLK1 | RT-PCR | B | forward: CCCTCACTGCCTTAACGACAAACC<br>reverse: GGCCGAGTTTGTACCACGTG | 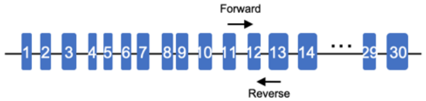 | NM_001110000 |
| | | O | forward: CCTCACGTGGTACAAACTCGGC<br>reverse: GTGGAGAGGGATTCCCAGATG | 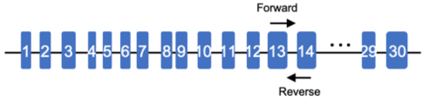 | NM_001278565 |
| SOX17 | RT-PCR | B | forward: GAGTCGCTGAGTCCCCTCG<br>reverse: CGCTTTAGCCGCTTCACCTG | 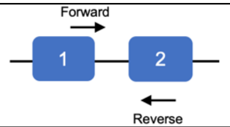 | NM_001206251 |
| | | O | forward: TGCACATGTACTACGGCCCC<br>reverse: TCACACGTCGGGATAGTTGCAG | 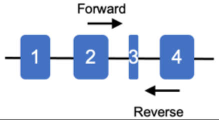 | XM_027972959 |
| PHEX | PCR | B | forward: CCAGAAGGCTTTTAATTGTCCATCC<br>reverse: GGACTTACAGATACAGCCTTGTC | 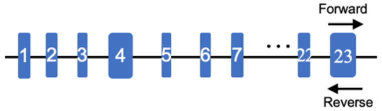 | NC_037357 |
| | | O | forward: CAATTCTACAATGAACAGAGGCGC | | NC_056808 |

| | | | | | |
|---|---|---|---|---|---|
| | | | reverse: GGGGCTTACAGATACAGCCTTGTC | 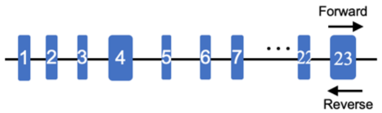 | |
| ZFY | PCR | 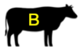 | forward: TGGCTCCTTTTTCCTTATGCACCAT | 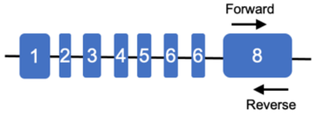 | NC_082638 |
| | | | reverse: TAATTATTGGTCCTGATGGACATCCC | | |
| | | 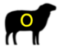 | forward: GATGGACATCCCTTGACTGTCTATCC | 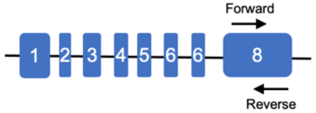 | NC_082741 |
| | | | reverse: CCCTTGTTCAGCTGTCTCATACTC | | |
| MSTN | PCR | 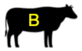 | forward: CATAAGCAAAATGATTAGTTTCTTT | 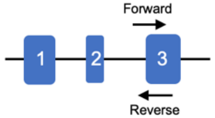 | NC_037329 |
| | | | reverse: AGCTTGTGCTTAAGTGACTG | | |
| | | 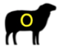 | forward: TCCATATGCAAATGGTTAGATGG | 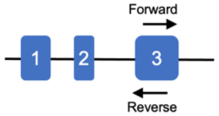 | NC_056055 |
| | | | reverse: TTGATTAACAAAATCCTAATTTACA | | |
| Episomal PE vector (Ampicillin) | Real-time PCR | — | forward: CTGCAACTTTATCCGCCTCC | — | — |
| | | | reverse: TCTGACAACGATCGGAGGAC | | |
| Actin | Real-time PCR | 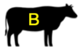 | forward: CCATCGGCAATGAGCGGTTCC | 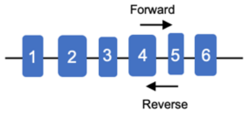 | NM_173979 |
| | | | reverse: CCGTGTTGGCGTAGAGGT | | |
| GAPDH | Real-time PCR | 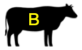 | forward: CATTGACCTTCACTACATGG | 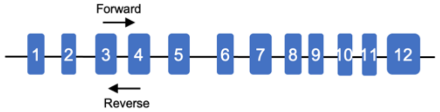 | NM_001034034 |
| | | | reverse: ATTGATGACGAGCTTCCCGT | | |
| IGF2/H19-DMR | Bisulfite PCR | 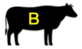 | forward: TTTTGTGGATTATTGTGGTATT | — | NC_037356 |
| | | | reverse: ATCTTAAACTAATCTCCCAACCC | | |

| | | | | | |
|---|---|---|---|---|---|
| | | O | forward: ATTTTAAATAGGGTTGAGAGGTTGT<br>reverse: AAACACAAAAAATCCCTCATTATC | — | NC_056074 |
| PEG10-DMR | Bisulfite PCR | B | forward: GTTTGGTATAGGTGTGGGATTT<br>reverse: TCAAAACCCTAAAAACTTAAATTCTC | — | NC_03733 |
| | | O | forward: GTTTGGTATAGGTGTGGGATT<br>reverse: TCTTCACACAATTTTAAAATACAACC | — | NC_056057 |
| PRDM1 | sgRNA | B | forward: CACCCTCGAGGACTTACGACTTGGCGAGTCA<br>reverse: AAACTGACTCGCCAAGTCGTAAGTCCTCGAG | 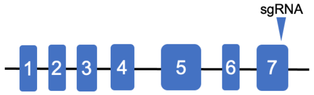 | NC_037336 |
| TFAP2C | sgRNA | B | forward: CACCCTCGAGGACTCCCAGATCATCTCGACC<br>reverse: AAACGGTCGAGATGATCTGGGAGTCCTCGAG | 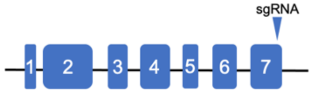 | NC_037340 |
| PRDM1 | PCR | B | forward: GCACAGTGGAGAACGACCTTTC<br>reverse: CCAGACCTCACCCATACAAAACC | 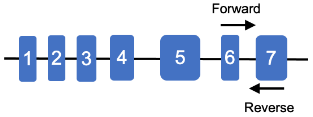 | NC_037336 |
| TFAP2C | PCR | B | forward: GGCTCGGGACTTTGCCTATG<br>reverse: GAGTCAAGAACGGAGCCGAAG | 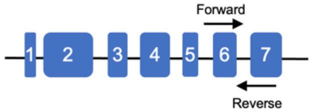 | NC_037340 |

**Table 7. List of pegRNA used in this study.**

| Target | Species | Spacer sequence | RTT sequence | PBS sequence |
|---|---|---|---|---|
| MSTN | 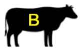 | ACGACAGCATCGAGATTCTG | TGTGACAG | AATCTCGATGCTGT |
| | 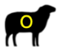 | ATTTATAAGTATTAAAATAA | AACATTCCATTA | TTTTAATACTTATAAAT |

**Table 8. MIQE checklist.**

| ITEM TO CHECK | IMPORTANCE | CHECKLIST | COMMENTS/WHERE |
|---|---|---|---|
| Definition of experimental and control groups | **E** | **YES** | Results and Methods |
| Number within each group | **E** | **YES** | Results and Methods |
| Assay carried out by core lab or investigator's lab? | D | **YES** | Investigator's lab |
| Acknowledgement of authors' contributions | D | **YES** | Author contributions |
| Description | **E** | **N/A** | |
| Volume/mass of sample processed | D | **N/A** | |
| Microdissection or macrodissection | **E** | **N/A** | |
| Processing procedure | **E** | **N/A** | |
| If frozen - how and how quickly? | **E** | **N/A** | |
| If fixed - with what, how quickly? | **E** | **N/A** | |
| Sample storage conditions and duration (especially for FFPE samples) | **E** | **N/A** | |
| Procedure and/or instrumentation | **E** | **YES** | Methods |
| Name of kit and details of any modifications | **E** | **YES** | Methods |
| Source of additional reagents used | D | **N/A** | |
| Details of DNase or RNAse treatment | **E** | **YES** | Methods |
| Contamination assessment (DNA or RNA) | **E** | **YES** | |
| Nucleic acid quantification | **E** | **YES** | Methods |
| Instrument and method | **E** | **YES** | Methods |
| Purity (A260/A280) | D | **YES** | Nanodrop (Thermal) and Supplementary Fig. 7 |
| Yield | D | **YES** | Supplementary Fig. 7 |
| RNA integrity method/instrument | **E** | **YES** | |
| RIN/RQI or Cq of 3' and 5' transcripts | **E** | **NO** | |

| | | | |
|---|---|---|---|
| Electrophoresis traces | D | **YES** | Denaturing agarose gel electrophoresis |
| Inhibition testing (Cq dilutions, spike or other) | **E** | **YES** | |
| Complete reaction conditions | **E** | **YES** | Methods |
| Amount of RNA and reaction volume | **E** | **YES** | Methods |
| Priming oligonucleotide (if using GSP) and concentration | **E** | **YES** | Methods |
| Reverse transcriptase and concentration | **E** | **YES** | Methods |
| Temperature and time | **E** | **YES** | Methods |
| Manufacturer of reagents and catalogue numbers | D | **YES** | Methods |
| Cqs with and without RT | D* | **N/A** | |
| Storage conditions of cDNA | D | **YES** | -20 °C |
| If multiplex, efficiency and LOD of each assay. | **E** | **YES** | Supplementary Fig. 7 |
| Sequence accession number | **E** | **YES** | Supplementary Table 6 |
| Location of amplicon | D | **YES** | Supplementary Table 6 |
| Amplicon length | **E** | **YES** | Results |
| *In silico* specificity screen (BLAST, etc) | **E** | **YES** | |
| Pseudogenes, retropseudogenes or other homologs? | D | **NO** | |
| Sequence alignment | D | **YES** | |
| Secondary structure analysis of amplicon | D | **NO** | |
| Location of each primer by exon or intron (if applicable) | **E** | **YES** | Supplementary Table 6 |
| What splice variants are targeted? | **E** | **N/A** | |
| Primer sequences | **E** | **YES** | Supplementary Table 6 |
| RTPrimerDB Identification Number | D | **N/A** | |
| Probe sequences | D** | **N/A** | |
| Location and identity of any modifications | **E** | **N/A** | |
| Manufacturer of oligonucleotides | D | **YES** | Beijing Genomics Institution (BGI) |

| | | | |
|---|---|---|---|
| Purification method | D | **YES** | HPLC |
| Complete reaction conditions | **E** | **YES** | Methods |
| Reaction volume and amount of cDNA/DNA | **E** | **YES** | Methods |
| Primer, (probe), Mg++ and dNTP concentrations | **E** | **YES** | Methods, Manufactures proprietory |
| Polymerase identity and concentration | **E** | **YES** | Methods |
| Buffer/kit identity and manufacturer | **E** | **YES** | Methods |
| Exact chemical constitution of the buffer | D | **NO** | Manufactures proprietory (Exact constitution of buffer not provided by the manufacturer) |
| Additives (SYBR Green I, DMSO, etc.) | **E** | **YES** | Methods (SYBR Green I) |
| Manufacturer of plates/tubes and catalog number | D | **YES** | 0.2 mL Polypropylene PCR Tube Strips and Flat Cap Strips (Axygen, PCR-0208-FCP-C) |
| Complete thermocycling parameters | **E** | **YES** | Methods |
| Reaction setup (manual/robotic) | D | **YES** | Manual setup |
| Manufacturer of qPCR instrument | **E** | **YES** | Methods |
| Evidence of optimisation (from gradients) | D | **YES** | Supplementary Fig. 7 |
| Specificity (gel, sequence, melt, or digest) | **E** | **YES** | Supplementary Fig. 7 |
| For SYBR Green I, Cq of the NTC | **E** | **YES** | Supplementary Fig. 7 |
| Standard curves with slope and y-intercept | **E** | **YES** | Supplementary Fig. 7 |
| PCR efficiency calculated from slope | **E** | **YES** | Supplementary Fig. 7 |
| Confidence interval for PCR efficiency or standard error | D | **NO** | |
| r2 of standard curve | **E** | **YES** | Supplementary Fig. 7 |
| Linear dynamic range | **E** | **YES** | Supplementary Fig. 7 (1 to 10 ng) |
| Cq variation at lower limit | **E** | **YES** | Supplementary Fig. 7 |
| Confidence intervals throughout range | D | **NO** | Supplementary Fig. 7 |

| | | | |
|---|---|---|---|
| Evidence for limit of detection | **E** | **YES** | Supplementary Fig. 7 (at no template control level) |
| If multiplex, efficiency and LOD of each assay. | **E** | **YES** | Supplementary Fig. 7 |
| qPCR analysis program (source, version) | **E** | **YES** | Methods |
| Cq method determination | **E** | **YES** | Methods |
| Outlier identification and disposition | **E** | **YES** | No outliers identified |
| Results of NTCs | **E** | **YES** | Supplementary Fig. 7 |
| Justification of number and choice of reference genes | **E** | **YES** | Methods |
| Description of normalisation method | **E** | **YES** | Methods |
| Number and concordance of biological replicates | D | **YES** | Supplementary Fig. 7 |
| Number and stage (RT or qPCR) of technical replicates | **E** | **YES** | Methods and Results |
| Repeatability (intra-assay variation) | **E** | **YES** | Methods and Results |
| Reproducibility (inter-assay variation, %CV) | D | **YES** | Methods and Results |
| Power analysis | D | **NO** | |
| Statistical methods for result significance | **E** | **YES** | Methods and Results |
| Software (source, version) | **E** | **YES** | Methods |
| Cq or raw data submission using RDML | D | **NO** | |

All essential information (E) must be submitted with the manuscript. Desirable information (D) should be submitted if available. If using primers obtained from RTPrimerDB, information on qPCR target, oligonucleotides, protocols and validation is available from that source.

*: Assessing the absence of DNA using a no RT assay is essential when first extracting RNA. Once the sample has been validated as DNA-free, inclusion of a no-RT control is desirable, but no longer essential.
**: Disclosure of the probe sequence is highly desirable and strongly encouraged. However, since not all commercial pre-designed assay vendors provide this information, it cannot be an essential requirement. Use of such assays is advised against.
